# Chronic IL-1 exposure drives LNCaP cells to evolve androgen and AR independence

**DOI:** 10.1101/2020.04.21.054452

**Authors:** H.C. Dahl, M. Kanchwala, S.E. Thomas-Jardin, A. Sandhu, P. Kanumuri, A.F. Nawas, C. Xing, C. Lin, D.E. Frigo, N.A. Delk

## Abstract

Chronic inflammation promotes prostate cancer (PCa) initiation and progression. We previously reported that acute intereluekin-1 (IL-1) exposure represses androgen receptor (AR) accumulation and activity, providing a possible mechanism for IL-1-mediated development of androgen- and AR-independent PCa. Given that acute inflammation is quickly resolved, and chronic inflammation is, instead, co-opted by cancer cells to promote tumorigenicity, we set out to determine if chronic IL-1 exposure leads to similar repression of AR and AR activity observed for acute IL-1 exposure and to determine if chronic IL-1 exposure selects for androgen- and AR- independent PCa cells. We generated isogenic sublines from LNCaP cells chronically exposed to IL-1α or IL-1β. Cells were treated with IL-1α, IL-1β, TNFα or HS-5 bone marrow stromal cells conditioned medium to assess cell viability in the presence of cytotoxic inflammatory cytokines. Cell viability was also assessed following serum starvation, *AR* siRNA silencing and enzalutamide treatment. Finally, RNA sequencing was performed for the IL-1 sublines. MTT, RT-qPCR and western blot analysis show that the sublines evolved resistance to inflammation-induced cytotoxicity and intracellular signaling and evolved reduced sensitivity to siRNA-mediated loss of *AR*, serum deprivation and enzalutamide. Differential gene expression reveals that canonical AR signaling is aberrant in the IL-1 sublines, where the cells show constitutive *PSA* repression and basally high *KLK2* and *NKX3.1* mRNA levels and bioinformatics analysis predicts that pro-survival and pro-tumorigenic pathways are activated in the sublines. Our data provide evidence that chronic IL-1 exposure promotes PCa cell androgen and AR independence and, thus, supports CRPCa development.

## Introduction

The tumor microenvironment is rich in inflammatory cytokines due to infiltrating immune cell paracrine secretion and tumor cell autocrine signaling[1, 2]. Normally, inflammatory cytokines signal the destruction and removal of foreign and damaged cells in wound healing[1, 2]. However, tumor cells can usurp inflammatory cytokines to promote tumor cell survival and disease progression[1, 2]. Inflammation results from pathogen infection, diet, or tissue injury, but if left unresolved, acute inflammation evolves into chronic inflammation, which drives prostate cancer (PCa) initiation and progression[3].

We and others have shown that acute treatment with the inflammatory cytokine, interleukin 1 (IL-1) represses androgen receptor (AR) accumulation and activity in AR^+^ PCa cell lines[4–7]. AR^+^ luminal cells form the bulk of the primary PCa tumor and require AR transcriptional activity for survival and proliferation[8]; thus, PCa therapies block androgen production (androgen deprivation therapy, ADT) or directly inhibit AR activity (anti-androgens)[8]. Interestingly, while acute IL-1 represses AR accumulation and activity in PCa cell lines, a subpopulation of the PCa cells still remain viable. Our RNA sequencing analysis of acute IL-1-treated PCa cell lines reveal that, along with repressing *AR* mRNA levels and AR signaling, IL-1 concomitantly upregulates pro-survival and tumorigenic molecules and pathways[7, 9]. These data suggest that IL-1 contributes to androgen and AR independence by selecting for PCa cells that remain viable independent of *AR* expression and/or AR activity.

In addition to pathogen infection, diet, or tissue injury, inflammation can also be induced by androgen deprivation[10–12]. Importantly, androgen deprivation-induced inflammation can lead to castration-resistant prostate cancer (CRPCa)[10–12]. For example, androgen deprivation induces PCa cells to secrete IL-1[12]. IL-1 recruits mesenchymal stem cells that secrete chemokine ligand 5 (CCL5) which promotes PCa cell stemness and castration resistance[12]. In addition to tumor cells, IL-1 also is produced by myeloid precursor cells, macrophages, and neutrophils in the tumor microenvironment[12, 13]; and our published data indicate that bone marrow stromal cell IL-1 paracrine signaling represses AR levels and activity in PCa cells[7]. Thus, IL-1 secreted by both tumor cells and infiltrating immune cells in the tumor microenvironment can contribute to PCa cell castration resistance.

Significantly, 10-20% of PCa patients will develop CRPCa and over 80% of CRPCa patients will have or develop incurable bone metastatic disease[14]. CRPCa typically emerges within 2 years following ADT[14], suggesting a time dependent evolution and selection for resistant cell populations. Thus, while acute (e.g. days) IL-1 exposure leads to repression of AR mRNA and protein levels in PCa cell lines[4–7], we sought to determine if chronic (e.g. months) IL-1 exposure would also select for cells that lose *AR* expression and evolve AR-independent survival.

In this study, we exposed the androgen-dependent, AR^+^ PCa cell line, LNCaP to IL-1 for several months and isolated IL-1 sublines. Surprisingly, while acute IL-1 exposure represses AR mRNA and protein accumulation, LNCaP cells chronically exposed to IL-1 restore AR and AR activity. However, the IL-1 sublines show enhanced viability in the presence of serum starvation, anti- androgen treatment and *AR* silencing. Thus, our data suggests that chronic IL-1 exposure indeed selects for castration-resistant PCa cells.

## Materials and Methods

### Cell Culture

LNCaP (ATCC, Manassas, VA; CRL-1740) and C4-2B (gift from Leland Chung) prostate cancer (PCa) cell lines were maintained in a 37°C, 5.0% (v/v) CO_2_ growth chamber, cultured in Dulbecco Modified Eagle Medium (DMEM (Gibco/Thermo Scientific; Grand Island, NY;1185-092) supplemented with 10% (v/v) fetal bovine essence (FB Essence (FBE); Seradigm, Radnor, PA; 3100-500), 0.4mM L-glutamine (L-glut; Gibco/Invitrogen, Paisley, PA; 25030-081), and

10U/ml penicillin G sodium and 10 mg/ml streptomycin sulfate (pen-strep; Gibco/ Invitrogen, Grand Island, NY; 15140-122).

### Chronic IL-1 Subline Generation and Maintenance

_L_NCaP cells were maintained in DMEM/10% FB Essence (FBE) containing 0.5 ng/ml IL-1α (Gold Bio, St. Louis, MO; 1110-01A-10) or IL-1β (Gold Bio, St. Louis, MO; 1110-01B-10) for at least 3 months. We chose 0.5 ng/ml concentration because our previous IL-1 dose response results demonstrate that LNCaP cells show the same molecular response in 0.5-25 ng/ml IL-1[15] and because IL-1 is cytotoxic, we chose the lowest dose to ensure we would be able to obtain surviving colonies. Thus, while IL-1 is cytotoxic and cytostatic for LNCaP cells[5, 15], after at least 3 months in 0.5 ng/ml IL-1, proliferative colonies emerge (after 3 months for cells in IL-1α and after 4 months for cells in IL-1β). The proliferative colonies were expanded and termed <u>LN</u>CaP IL-1<u>α</u> <u>s</u>ubline (LNas) and <u>LN</u>CaP IL-1<u>β</u> <u>s</u>ubline (LNbs). We generated three LNas sublines and two LNbs sublines which all show similar phenotypes; therefore, LNas1 and LNbs1 are herein characterized. During subline generation, LNCaP parental cells were cultured in vehicle control (1 x phosphate buffer saline, PBS) (Corning, Manassas, VA; 21-040-CM) alongside the sublines. During the 3-4-month period before the sublines began to proliferate in IL-1, we made frozen stocks of the proliferating LNCaP parental cells. Once the sublines began to proliferate in IL-1 and, thus, could be expanded for experiments, LNCaP parental cells of similar passage number were brought back up from the frozen stocks to be characterized alongside the IL-1 sublines. Cell line authentication for LNCaP, LNas1, LNbs1 was performed by STR profiling by the DNA Genotyping Core, UTSW Medical Center. All cell lines used in the study are commercially available and purchased from American Tissue Culture Collection (ATCC). We hereby confirm that none of the used cell lines require any ethics approval for their use.

### Cell Treatments

#### Cytokines

Human recombinant IL-1α (GoldBio, St. Louis, MO; 1110-01A-100), IL-1β (GoldBio, St. Louis, MO; 1110-01B-100), or TNFα (Goldbio, St. Louis, MO; 1130-01-10) were resuspended in 0.1% bovine serum albumin (BSA) (Thermo Fisher Scientific, Fair Lawn, NJ; BP 1600-1) in 1X phosphate buffered saline (PBS; Corning, Manassas, VA; 21-040-CM). Cells were treated with vehicle control (0.1% BSA in 1X PBS), IL-1 or TNFα added to DMEM/10% FBE growth medium. *<u>HS-5 condition media (CM):</u>* HS-5 bone marrow stromal cells were grown in DMEM/10% FBE for 5 days, the medium collected and filtered (EMD Millipore, Burlington, MA; SCGPU05RE; pore size 0.220μm) to remove cell debris. *<u>Interleukin-1 Receptor Antagonist (IL-1RA):</u>* Cells were pre-treated for 1 day with vehicle control (0.1% BSA in 1X PBS) or 400 ng/ml human recombinant IL-1RA (R&D Systems, Minneapolis, MN; 280-RA/CF) 0 in the DMEM/10% FBE growth medium. The next day, the medium was removed and replaced with fresh DMEM/10% FBE or HS- 5 CM, plus an additional 400 ng/ml IL-1RA or vehicle 0 control, for 3 days. Gene silencing (siRNA): The following siRNA concentrations were used: 70nM non-targeting siRNA (Dharmacon, Lafayette, CO; D-001206-13-05) or *AR* siRNA (Dharmacon, Lafayette, CO; M-003400-02-0005, pool of 4 oligos). Cells were transfected with siRNA using siTran 1.0 transfection reagent (Origene, Rockville, MD; TT300003) DMEM/2.5% FBE growth medium for 4 days. *<u>Serum starvation</u>*: Cells were plated in DMEM/2.5% FBE growth medium and once the cells were attached and semi-confluent, the medium was replaced with DMEM/0% FBE (serum starvation) or DMEM/10% FBE (replete medium control). *<u>R1881:</u>* Cells were plated and maintained in DMEM/2.5% FBE growth medium and once the cells were attached and semi-confluent, the medium was replaced with DMEM/0% FBE plus 10 nM R1881 (Sigma-Aldrich, St. Louis, MO; R0908) or vehicle control (Dimethyl sulfoxide, DMSO; Corning, Manassas, VA; 25-950-CQC) for 4 days. *<u>Enzalutamide (ENZA):</u>*Cells were plated and maintained in DMEM/2.5% FBE growth medium and once the cells were attached and semi-confluent, the medium was replaced with DMEM/2.5% FBE plus 50 μM ENZA (Selleckchem, Houston, TX; S1250) or vehicle control (DMSO) for 3 or 5 days.

### RNA Isolation and Reverse Transcription Quantitative PCR (RT-qPCR)

Total RNA was extracted, reverse transcribed, and analyzed by RT-qPCR as previously described[7]. Primer sequences for genes of interest are listed below. Gene of interest cycle times (CT) were normalized to the β-actin. Relative mRNA levels were calculated using the 2-β CT method. Primer sequences, 5’-3’: Androgen Receptor (AR), forward ΔΔ AAGACGCTTCTACCAGCTCACCAA, reverse TCCCAGAAAGGATCTTGGGCACTT; Beta actin (β-actin), forward GATGAGATTGGCATGGCT TT, reverse CACCTTCACCGGTCCAGTTT; β *Prostate Specific Antigen (PSA)*, forward CACCTGCTCGGGTGATTCTG, reverse ACTGCCCCATGACGTGATAC; *Mitochondrial Superoxide Dismutase 2 (SOD2)*, forward GGCCTACGTGAACAACCTGA, reverse GTTCTCCACCACCGTTAGGG; *RELA*, forward TGAACCAGGGCATACCTGTG, reverse CCCCTGTCACTAGGCGAGTT; *NFKB1*, forward TGATCCATATTTGGGAAGGCCTGA, reverse GTATGGGCCATCTGTTGGCAG; Kallikrein-2 (*KLK2)*, forward AGCCTCCATCTCCTGTCCAA, reverse CCCAGAATCACCCCCACAAG; *NK3 homeobox 1 (NKX3.1)*, forward CCCACACTCAGGTGATCGAG, reverse GTCTCCGTGAGCTTGAGGTT; *Interleukin-1 Receptor Type 1(IL1R1)*, forward TGGGGAAGGGTCTACCTCTG, reverse TCCCCAACGTAGTCATCCCT; *MAPK11* forward CGACGAGCACGTTCAATTCC, reverse TCACAGTCCTCGTTCACAGC; *MAPK13* forward CTGCCCAAGACCTACGTGTC, reverse GGTCGGCTCAGCTTCTTGAT; *MAPK8* forward CTCGCTACTACAGAGCACCC, reverse CTCCCATAATGCACCCCACA; *CREBBP* forward TGAGAACTTGCTGGACGGAC, reverse GCTGTCATTCGCCGAGAAAC; *FZD1* forward CATCGTCATCGCCTGCTACT, reverse TAGCGTAGCTCTTGCAGCTC; *FZD3* forward TGGCCCTTGACTGTGTTCAT, reverse CAAAGCTGCTGTCTGTTGGTC; *FZD4* forward GCAGGACTCAAATGGGGTCA, reverse CAATGGTTTTCACTGCGGGG; *FZD9* forward AAGATCATGAAGACGGGCGG, reverse TAGCAAACGATGACGCAGGT.

### RNA-Sequencing (RNA-seq) Analysis

RNA-seq was performed by the Genome Center at the University of Texas at Dallas (Richardson, TX). Total RNA library was prepared using Illumina Truseq Stranded Total RNA prep Gold kit (Illumina). The prepared libraries were sequenced on an Illumina NextSeq 500 platform (San Diego, CA) with 75bp single-end reads. Fastq files were checked for quality using fastqc (v0.11.2)[16] and fastq_screen (v0.4.4)[17] and were quality trimmed using fastq-mcf (ea-utils/1.1.2-806)[18]. Trimmed fastq files were mapped to hg19 (UCSC version from igenomes) using TopHat[19], duplicates were marked using picard-tools (v1.127 https://broadinstitute.github.io/picard/), read counts were generated using featureCounts[20] and differential expression analysis was performed using edgeR[21] and limma[22]. Differential gene expression lists were generated using the following cut-offs: log_2_ counts per million (CPM) ≥ 0, log_2_ transcripts per million (TPM) ≥ 0, log_2_ fold change (FC) ≥ 0.6 or ≤ -0.6, false discovery rate (FDR) ≤ 0.05, adjusted p-value ≤ 0.05. Pathway analysis was conducted using QIAGEN’s Ingenuity Pathway Analysis (IPA) tool (http://www.qiagen.com/ingenuity) RNA-seq datasets generated for this study are available at GEO NCBI, accession GSE142706. RNA-seq dataset generated by Poluri et al. is available at GEO NCBI, accession GSE128749.

### Western Blot

Protein was isolated from cells using NP40 lysis buffer (0.5% NP40 [US Biological, Salem, MA; N3500], 50 mM of Tris [pH 7.5], 150 mM of NaCl, 3 mM of MgCl2, 1X protease inhibitors [Roche, Mannheim, Germany; 05892953001]). Protein concentration was measured using the Pierce BCA Protein Assay Kit (Thermo Fisher Scientific, Waltham, MA; 23227). For Western blot analysis, equal protein concentrations were loaded onto and separated in 12% (wt/vol) sodium dodecyl sulfate polyacrylamide gel (40% acrylamide/bisacrylamide solution; Bio-Rad, Hercules, CA; 161‐0148). Proteins were transferred from the gel to 0.45 m pore size μ nitrocellulose membrane (Maine Manufacturing, Sanford, ME; 1215471) and total protein visualized using Ponceau S (Amresco, Radnor, PA; K793). The membrane was blocked with 2.5% (wt/vol) BSA (Thermo Fisher Scientific, Waltham, MA; BP 1600-1) in 1X tris-buffered saline with Tween 20 (TBST; 20 mM of Tris, pH 7.6, 150 mM of NaCl, 0.05% Tween-20). Primary and secondary antibodies were diluted in 2.5% BSA in 1X TBST. Protein blot bands were visualized using Clarity Western ECL Substrate (Bio-Rad, Hercules, CA; 1705061) or SuperSignal West Femto Maximum Sensitivity Substrate (Thermo Scientific, Rockford, IL; 34095) and imaged using Amersham Imager 600 (GE, Marlborough, MA). *<u>Primary antibodies:</u>* AR (Cell Signaling, Danvers, MA; D6F11), PARP (Cell Signaling, Danvers, MA; 9532S), SOD2 (Abgent, San Diego, CA; AM7579a), β-actin (Santa Cruz, Santa Cruz, CA; sc-69879), NFKB1β (Cell Signaling, Danvers, MA; 3035S), PSA (Cell Signaling, Danvers, MA; 5365S), RELA (Cell Signaling, Danvers, MA, L8F6) or NKX3.1(Cell Signaling, Danvers, MA, D2Y1A), Caspase 3 (Cell Signaling, Danvers, MA; 9664S), Cleaved Caspase 3 (Cell Signaling, Danvers, MA; 9664S). *<u>Secondary antibodies:</u>* Sheep anti-mouse (Jackson ImmunoResearch Laboratories, Grove, PA; 515-035-062), goat anti-rabbit (Abnova, Walnut, CA; PAB10822). Western blot densitometry was performed using Image J (National Institutes of Health, Bethesda, Maryland). β-actin or ponceau stain is the western blot loading control and the protein/β-actin or protein/ponceau stain ratio is normalized to treatment control for densitometry.

### MTT (3-(4,5-Dimethylthiazol-2-yl)-2,5-diphenyltetrazolium bromide) Viability Assay

MTT assay (Trevigen, Gaithersburg, MD; 4890-25-K) was performed according to manufacturer’s instructions. Cell viability was quantified as the optical density (OD) read at wavelengths 540 nm and 650 nm. The final OD was calculated as follows: OD 540 nm – OD 650 nm. OD was measured using the Cytation3 Imaging Reader (BioTek, Winooski, VT).

### Cell Counts

Growth medium was removed along with dead cell debris and the viable cells were fixed in 100% cold methanol. The nuclei were stained with DAPI (Roche Diagnostics, 10236276001) and the stained cells imaged and counted using the Cytation3 Cell Imaging Multi-Mode Reader (BioTek, Winooski, Vermont).

### Statistical Analysis

Statistical significance for MTT and cell counts was determined using unpaired student t test calculated using Microsoft Excel. P-values of ≤ 0.05 were considered to be statistically significant and denoted by asterisks(*p ≤ 0.05; **p ≤ 0.05***p ≤ 0.005). Graphs are shown as the average of ≤ a minimum of n = 3 biological replicates +/- standard deviation (STDEV).

## Results

### LNCaP cells chronically treated with IL-1 restore AR and AR activity

IL-1 encompasses a family of related cytokines. The two most biologically and clinically relevant IL-1 family members are IL-1 alpha (IL-1α) and IL-1 beta (IL-1β). IL-1α and IL-1β both signal through the IL-1 receptor (IL-1R1), they elicit both similar and unique cell responses,[13] and both IL-1α and IL-1β are clinically relevant in prostatic disease[23–25]. We previously showed that acute IL-1 treatment (e.g., days) represses AR levels and AR target gene expression in LNCaP cells[4, 7]. Therefore, we hypothesized that, likewise, chronic IL-1 exposure would repress AR accumulation and activity. We treated LNCaP cells with 12.5 ng/ml IL-1α or IL-1β for 3 days or 1, 2, or 3 weeks and determined AR mRNA or protein accumulation by RT-qPCR or western blot, respectively. IL-1 repressed AR mRNA (Fig. 1A) and protein (Fig. 1B) levels at day 3, however, AR mRNA and protein levels began to re-emerge at 1-2 weeks in IL-1-treated LNCaP cells. Thus, while acute IL-1 exposure represses AR mRNA and protein accumulation in LNCaP cells, our experiment shows that *AR* expression re-emerges in extended IL-1 treatment.

**Fig 1.**
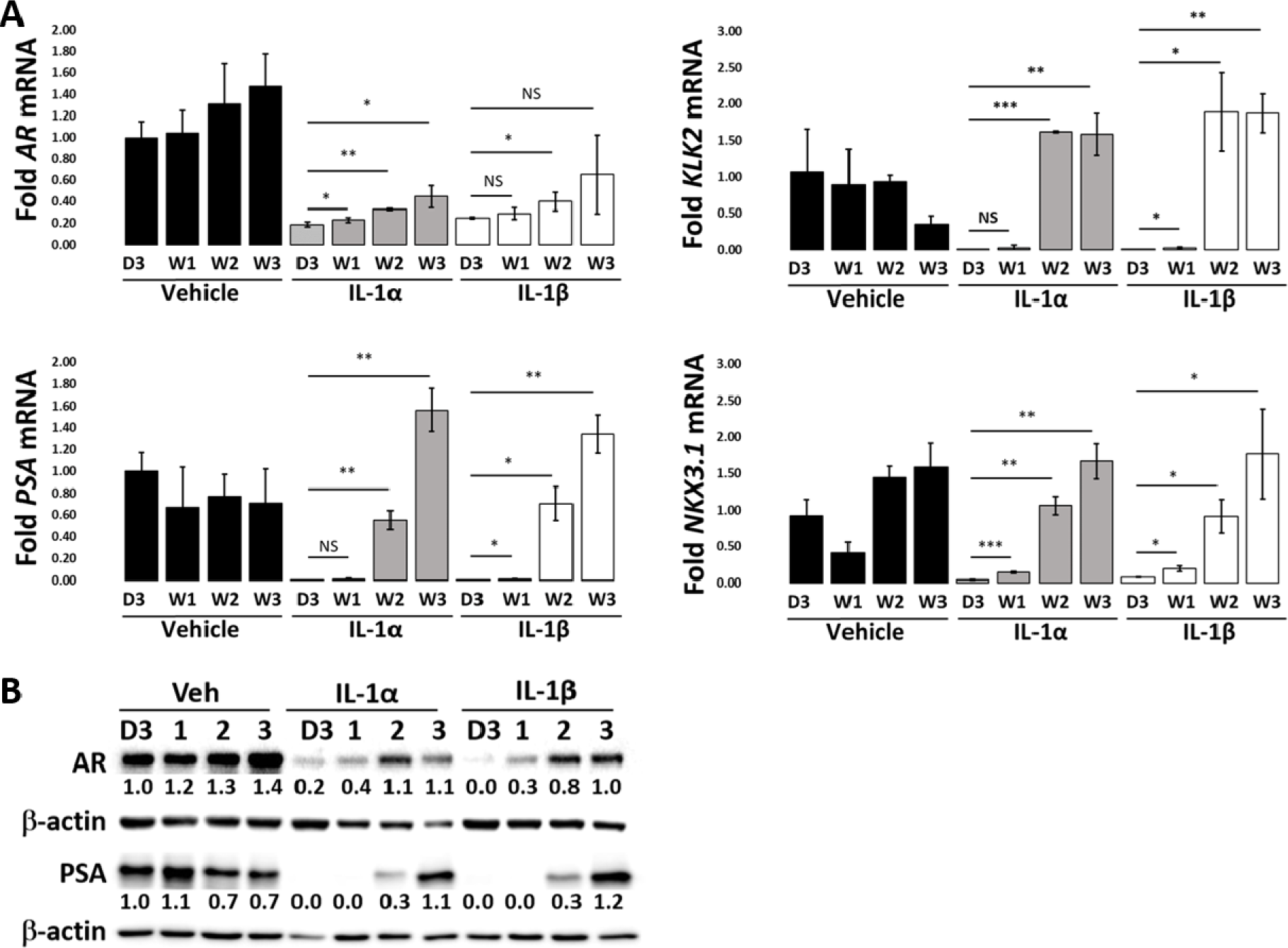
LNCaP cells lose sensitivity to IL-1-mediated AR repression over time. (A) RT- qPCR and (B) western blot analyses were performed for LNCaP cells treated with vehicle control, 12.5 ng/ml IL-1α or IL-1β for 3 days (D3) or 1 (W1), 2 (W2) or 3 (W3) weeks. mRNA and/or protein accumulation were determined for AR and AR target genes, *KLK2*, *PSA* and *NKX3.1*. IL-1 repressed (A) mRNA and (B) protein for AR and AR target genes, *KLK2*, *PSA* and *NKX3.1* at 3 days. AR and AR target gene levels began to re-emerge after 1-2 weeks of c0.05, ** 0.005, ≤ *** 0.0005. mRNA fold change is normalized to day 3 vehicle control. Western blot band densitometry is normalized to β-actin and fold change normalized to day 3 vehicle control.

To assay AR activity, we determined mRNA and/or protein accumulation of the AR target genes, *Kallikrein Related Peptidase 2* (*KLK2*), *Prostate Specific Antigen* (*PSA/KLK3*) and *NK3 Homeobox 1* (*NKX3.1*) in LNCaP cells treated with 12.5 ng/ml IL-1α or IL-1β for 3 days or 1, 2, or 3 weeks. IL-1 repressed *KLK2*, *PSA*, and *NKX3.1* mRNA (Fig. 1A) and PSA protein (Fig. 1B) levels at day 3; however, mRNA and proteins levels began to re-emerge at 1 to 2 weeks in IL-1- treated LNCaP cells. Our experiments show that LNCaP cells begin to lose sensitivity to IL-1- mediated repression of AR activity at 1 to 2 weeks of chronic exposure to IL-1. Thus, our experiments suggest that, while acute IL-1 exposure represses AR activity, PCa cells can evolve insensitivity to chronic IL-1 signaling.

### The generation of chronic IL-1 LNCaP sublines

Chronic inflammation, including IL-1β and IL-1 pathway signaling molecules, has been shown to promote castration resistance in Myc-CaP allograft and LNCaP xenograft models[10–12,26]. Given that castration-resistant PCa (CRPCa) cells can be AR-dependent[8] or AR- independent[27], we set out to determine if chronic IL-1α or IL-1β exposure promotes AR- dependent or AR-independent castration resistance in LNCaP cells. To address our question, we cultured LNCaP cells in 0.5 ng/ml IL-1α or IL-1β for 3-4 months. IL-1 is cytostatic and cytotoxic for LNCaP cells[5, 15] (Fig. 2A), but after 3-4 months of culturing in IL-1α or IL-1β, the surviving cells began to proliferate. We expanded the surviving cell subpopulations and designated the isolated cells as <u>LN</u>CaP IL-1<u>α</u> <u>s</u>ubline (LNas1) or <u>LN</u>CaP IL-1<u>β</u> <u>s</u>ubline (LNbs1).

**Fig 2.**
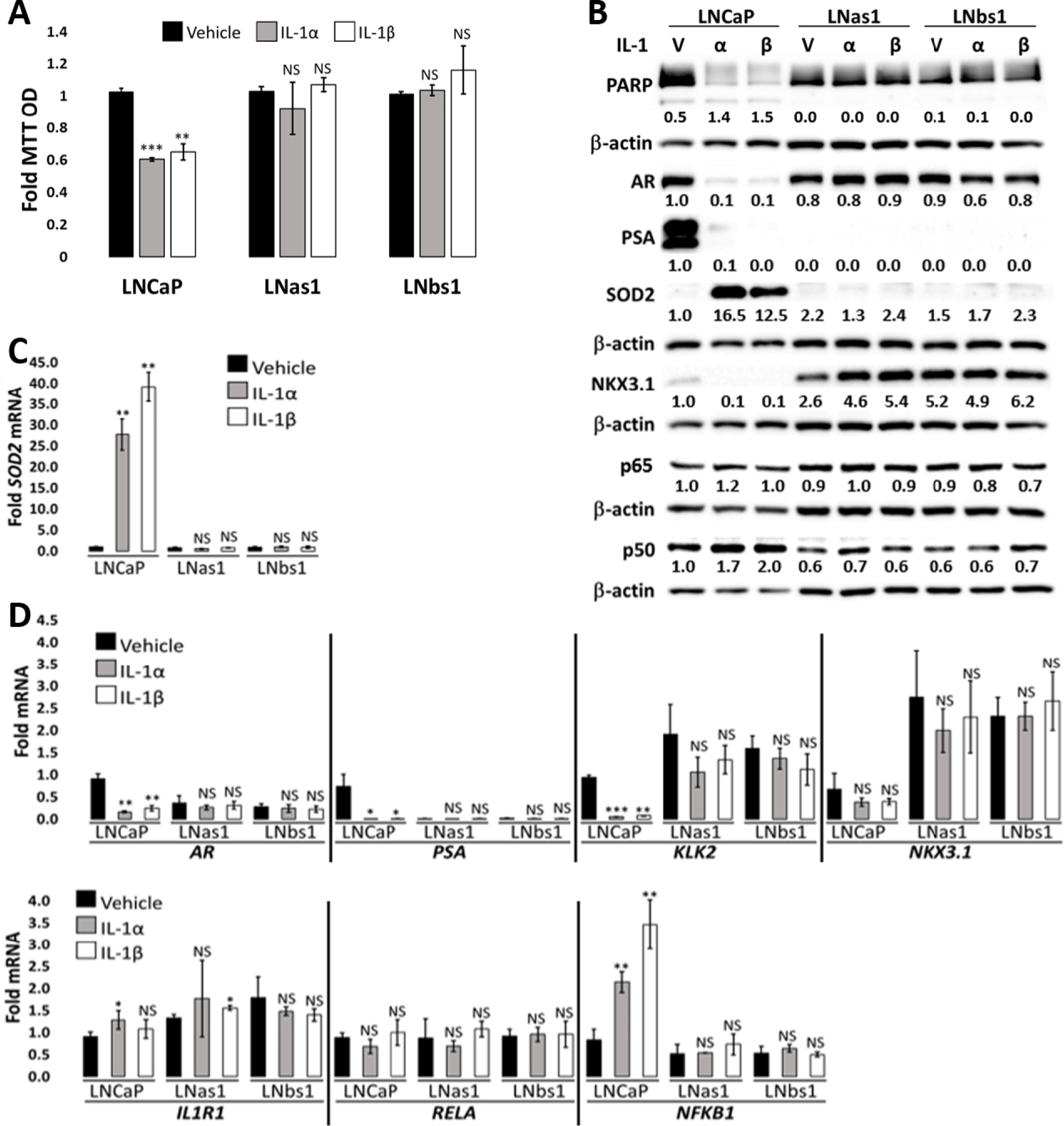
LNas1 and LNbs1 IL-1 sublines are resistant to IL-1-induced cytotoxicity and intracellular signaling. (A) Cell viability was determined using MTT for LNCaP, LNas1 and LNbs1 for 5 days with vehicle control, 25 ng/ml IL-1α or IL-1β. IL-1 reduces LNCaP cell viability; however, LNas1 and LNbs1 cells remain viable. (B) LNCaP, LNas1 and LNbs1 cells were treated for 3 days with vehicle control or 25 ng/ml IL-1α or IL-1β and analyzed for protein accumulation by western blot. Western blot analyses were performed for IL-1 target gene, SOD2, to determine treatment efficacy and activation of IL-1 intracellular signaling, PARP cleavage to determine activation of apoptosis, and AR, PSA, and NKX3.1 to determine AR activity. p65 and p50 are NFκB transcription factor subunits. IL-1 induced SOD2 protein accumulation and PARP cleavage in LNCaP cells, but IL-1 had less or no effect on SOD2 protein levels or PARP cleavage in LNas1 or LNbs1 cells. IL-1 downregulated AR, PSA and NKX3.1 protein accumulation in LNCaP cells, but IL-1 had less or no effect on the protein levels in LNas1 and LNbs1 cells. IL-1 had no effect on p65 protein levels in LNCaP, LNas1 or LNbs1. IL-1 induced p50 protein in LNCaP cells, but not in LNas1 or LNbs1 cells. (C, D) LNCaP, LNas1 and LNbs1 cells were treated for 3 days with vehicle control or 25 ng/ml IL-1α or IL-1β and analyzed for mRNA levels by RT-qPCR for *SOD2*, *AR*, *PSA*, *KLK2*, *NKX3.1*, *IL-1R1*, *RELA* (p65) and *NFKB1* (p50). IL-1 significantly induced *SOD2* and *NFKB1*, significantly repressed *AR*, *PSA*, *KLK2*, and slightly repressed *NKX3.1* mRNA levels in LNCaP cells, but IL-1 had no significant effect on the mRNA levels in LNas1 or LNbs1 cells. IL-1 had little or no effect on *IL-1R1* or *RELA* mRNA levels in LNCaP, LNas1 or LNbs1 cells. Finally, compared to LNCaP cells, LNas1 and LNbs1 have lower basal PSA and (p50) mRNA or protein levels and higher basal KLK2 and NKX3.1 mRNA or protein levels. Error bars, ± STDEV of 3 biological replicates; p-0.05, **≤0.005. For each individual cell line, fold MTT optical density (OD) is normalized to the vehicle control. Western blot band densitometry is normalized to β-actin and fold change normalized to LNCaP vehicle control. PARP cleavage densitometry shows the ratio of cleaved to uncleaved PARP. Fold mRNA levels are normalized to LNCaP vehicle control for IL-1 treatments in order to also compare basal levels between the cell lines.

### LNas1 and LNbs1 sublines are insensitive to IL-1 and TNFα cytotoxicity

While IL-1 is cytotoxic for LNCaP cells, by virtue of the selection of the sublines in chronic IL-1, the LNas1 and LNbs1 sublines are insensitive to IL-1-induced cell death. To demonstrate, LNCaP, LNas1 and LNbs1 cells were treated with 25 ng/ml IL-1α or IL-1β for 5 days and cell viability was determined using MTT. IL-1 reduced LNCaP cell viability by ∼40%, but IL-1 had no significant effect on LNas1 or LNbs1 cell viability (Fig. 2A). Apoptosis activation was assessed intracellularly by PARP and caspase 3 cleavage western blot in cells treated with 25 ng/ml IL-1α or IL-1β for 3 days. IL-1 induced PARP and caspase 3 cleavage in LNCaP cells but had little or no effect on PARP cleavage in LNas1 or LNbs1 (Fig. 2B, S1A Fig). Thus, the IL-1 sublines are insensitive to IL-1-induced cell death.

It was previously reported that secreted factors from the bone marrow stromal cell line, HS-5, induce apoptosis in LNCaP cells[28], where HS-5 cells secrete IL-1α and IL-1β, among other cytokines[29]. In kind, we found that IL-1 Receptor Antagonist (IL1RA) is sufficient to block PARP and caspase 3 cleavage in LNCaP cells exposed to conditioned medium (CM) from HS-5 cells (Fig. 3A, S1B Fig.), indicating that IL-1α or IL-1β are sufficient to mediate HS-5 CM-induced LNCaP apoptosis. We treated LNCaP, LNas1 and LNbs1 cells with HS-5 CM and analyzed cell viability using MTT on day 5 or assessed PARP and caspase 3 cleavage on day 3. As observed for IL-1 treatment, LNas1 and LNbs1 sublines are insensitive to HS-5 CM-induced cell death (Fig. 3B & C, S1C Fig.).

**Fig 3.**
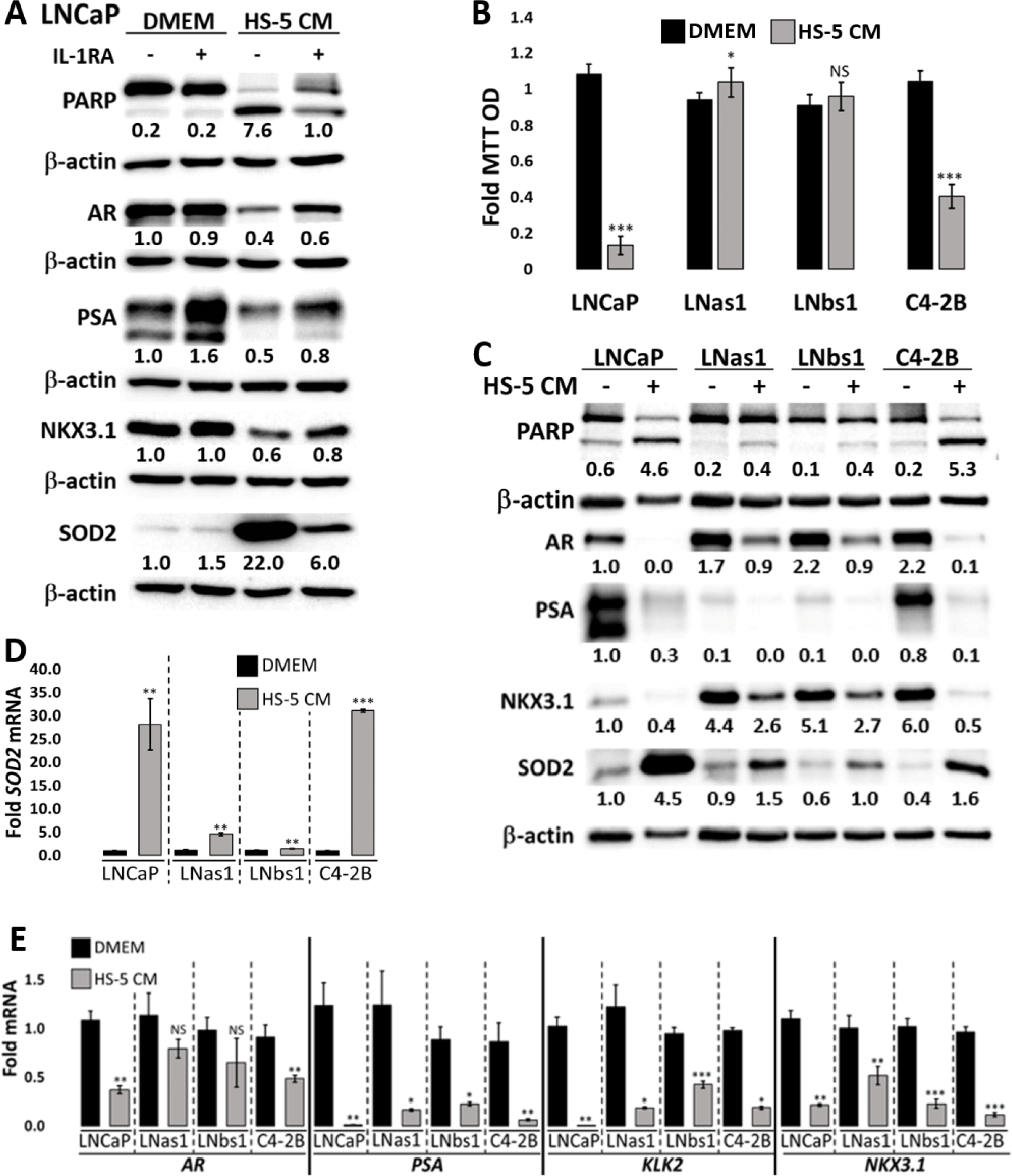
LNas1 and LNbs1 IL-1 sublines are resistant to HS-5-induced cytotoxicity and intracellular signaling. (A) LNCaP cells were pre-treated for 1 day with vehicle control or 400 ng/ml human recombinant IL-1RA and the following day the medium was replaced with treatment control (DMEM) or HS-5 conditioned medium (CM) plus an additional 400 ng/ml IL-1RA or vehicle control for 3 additional days. Western blot analyses were performed for SOD2 to determine treatment efficacy, PARP cleavage to determine activation of apoptosis, and AR, PSA, and NKX3.1 to determine AR activity. IL-1RA was sufficient to attenuate HS-5 CM- induced SOD2 upregulation, PARP cleavage, and repression of AR, PSA and NKX3.1. (B) Cell viability was determined using MTT for LNCaP, LNas1, LNbs1 and C4-2B cells treated for 5 days with treatment control or HS-5 CM. HS-5 CM reduced LNCaP and C4-2B cell viability; however, LNas1 and LNbs1 cells remained viable. LNCaP, LNas1, LNbs1 and C4-2B cells treated for 3 days with treatment control or HS-5 CM and analyzed for protein accumulation by western blot (C) or mRNA levels by RT-qPCR (D, E). Protein and/or RNA analyses were performed for IL-1 target gene, *SOD2*, to determine treatment efficacy and activation of IL-1 intracellular signaling, PARP cleavage to determine activation of apoptosis, and AR, PSA, KLK2 and NKX3.1 to determine AR activity. HS-5 CM induced PARP cleavage and SOD2 mRNA and protein levels and repressed AR, PSA, KLK2 and NKX3.1 mRNA and protein levels in LNCaP and C4-2B cells, but HS-5 CM had less effect in LNas1 or LNbs1 cells. Error bars, ± STDEV of 0.005, ***≤0.0005. For each individual cell line, fold MTT optical density (OD) is normalized to the treatment control (DMEM). Western blot band densitometry is normalized to β-actin and fold change normalized to LNCaP treatment control (DMEM). PARP cleavage densitometry shows the ratio of cleaved to uncleaved PARP. For each individual cell line, fold mRNA levels are normalized to the cell line treatment control (DMEM); basal levels are not compared between the cell lines.

Like IL-1, Tumor Necrosis Factor alpha (TNFα) is also a cytotoxic inflammatory cytokine that likewise signals through the Nuclear Factor kappa B (NFκB) intracellular signaling pathway[30]. Therefore, we determined if LNas1 and LNbs1 cells are also insensitive to TNFα-induced cytotoxicity. We treated LNCaP, LNas1 and LNbs1 cells with 25 ng/ml TNFα for 5 days and analyzed cell viability using MTT. TNFα induced an ∼80% reduction in LNCaP cell viability, but TNFα had no significant effect on LNas1 or LNbs1 cell viability (Fig. 4A). Taken together, chronic IL-1 exposure selects for LNCaP cells that evolve resistance to inflammatory cytokine cytotoxicity.

**Fig 4.**
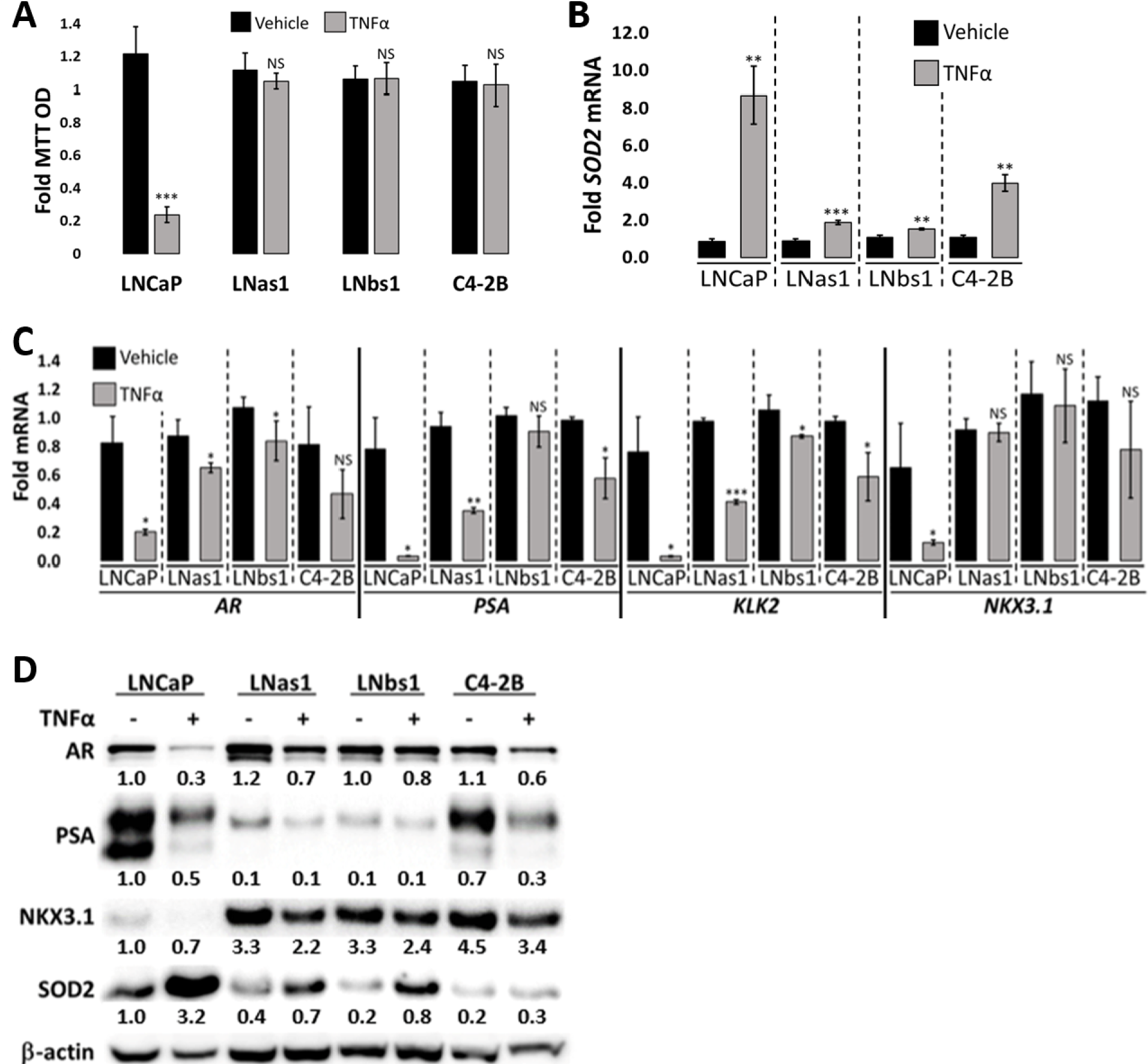
LNas1 and LNbs1 IL-1 sublines are resistant to TNFα-induced cytotoxicity and intracellular signaling. (A) Cell viability was determined using MTT for LNCaP, LNas1, LNbs1 and C4-2B cells treated for 5 days with vehicle control or 25 ng/ml TNFα. TNFα reduced LNCaP cell viability; however, LNas1, LNbs1 and C4-2B cells remain viable. LNCaP, LNas1, LNbs1 and C4-2B cells were treated for 3 days with treatment control or 25 ng/ml TNFα and analyzed for mRNA levels by RT-qPCR (B, C) or protein accumulation by western blot (D). RNA and/or protein analyses were performed for TNFα target gene, *SOD2*, to determine treatment efficacy and activation of TNFα intracellular signaling and for AR, PSA, KLK2 and NKX3.1 to determine AR activity. TNFα induced SOD2 and repressed AR, PSA, KLK2 and NKX3.1 mRNA and protein levels in LNCaP cells, but TNFα had less or no effect on mRNA or protein levels in LNas1, LNbs1 or C4-2B cells. Error bars, ± STDEV of 3 biological replicates; p-value, *≤0.05, ≤0.005, *** 0.0005. For each individual cell line, fold MTT optical density (OD) is normalized to the vehicle control. For each individual cell line, fold mRNA levels are normalized to the cell line vehicle control; basal levels are not compared between the cell lines. Western blot band densitometry is normalized to β-actin and fold change normalized to LNCaP vehicle control.

### Insensitivity to IL-1 or TNFα cytotoxicity is characteristic of the castration-resistant LNCaP subline, C4-2B

We previously reported that the castration-resistant LNCaP subline, C4-2B, shows reduced sensitivity to IL-1-induced cytotoxicity[15]. Interestingly, similar to LNas1 and LNbs1 cells, C4-2B also showed reduced sensitivity to TNFα-induced cytotoxicity (Fig. 4A). C4-2B cells are also slightly less sensitive to HS-5 CM-induced cytotoxicity (∼60% reduced cell viability) as compared to LNCaP cells (∼85% reduced cell viability), but C4-2B cells are not as resistant as LNas1 or LNbs1 cells (Fig. 3B). Indicative of C4-2B sensitivity to HS-5 CM, PARP and caspase 3 cleavage is also induced in the C4-2B cells (Fig. 3C, S1D Fig.). These results suggest that, like chronic exposure to inflammation, androgen deprivation also selects for cells that evolve resistance to inflammatory cytokine cytotoxicity; but not surprisingly, resistance is more pronounced in cells that are selected through specific exposure to chronic inflammation.

### LNas1 and LNbs1 sublines lose IL-1 and TNFα intracellular signaling responses

IL-1 and TNFα bind their respective receptors to activate the NFκB transcription factor and NFκB induces transcription of genes that mediate cell response to inflammatory cytokine signaling[1, 31]. Having observed that the LNas1 and LNbs1 sublines remain viable in the presence of the cytotoxic inflammatory cytokines, IL-1α, IL-1β, TNFα and HS-5 CM (Fig. 2A, 3B & 4A), we wanted to determine if canonical NF-κB signaling is disrupted in LNas1 and LNbs1 cells. *Super Oxide Dismutase 2* (*SOD2*) is a canonical NFκB-induced gene upregulated in response to IL-1 or TNFα exposure[32]. HS-5 CM also induces SOD2 accumulation and IL-1RA is sufficient to attenuate SOD2 induction in HS-5 CM-treated LNCaP cells[7] (Fig. 3A), indicating that IL-1α or IL-1β are sufficient to mediate HS-5 CM-induced SOD2 upregulation.

We treated LNCaP, LNas1, LNbs1 and C4-2B cells with 25 ng/ml IL-1α, IL-1β or TNFα, or HS-5 CM for 3 days and determined SOD2 mRNA and protein levels by RT-qPCR or western blot, respectively. IL-1 (Fig. 2B & C), HS-5 CM (Fig. 3C & D) and TNFα (Fig. 4B & D) increased SOD2 mRNA and protein accumulation in LNCaP cells, but, comparatively, IL-1, HS-5 CM and TNFα had lesser or no detectable effect on SOD2 accumulation in LNas1 and LNbs1 sublines. Likewise, IL-1[15] and TNFα (Fig. 4B & D) had less of an effect on SOD2 induction in C4-2B cells than in LNCaP cells. C4-2B cells, however, did show a similar *SOD2* mRNA induction response to HS-5 CM as the LNCaP cells (Fig. 3D), which is likely due to a synergistic effect of the milieu of cytokines presents in HS-5 CM. Notwithstanding, HS-5 CM-induced SOD2 protein accumulation was lower in C4-2B cells than in LNCaP cells (Fig. 3C). Thus, LNas1, LNbs1 and C4-2B cells show no or reduced canonical intracellular response to IL-1, TNFα or HS-5 CM.

Finally, in addition to SOD2 induction, IL-1 (Fig. 2B & D), HS-5 CM (Fig. 3C & E) and TNFα (Fig.4C & D) repress AR and AR target gene accumulation in LNCaP cells[4–7,15,30,33]; and we and others have shown that IL-1[15] or TNFα[30, 33] can repress *AR* mRNA levels and AR activity in a NF-κB-dependent manner. As compared to LNCaP cells, LNas1 and LNbs1 cells showed lesser or no repression of AR or AR target genes, *PSA*, *KLK2* or *NKX3.1*, in response to IL-1 (Fig. 2B & D), HS-5 CM (Fig. 3C & E) and TNFα (Fig. 4C & D). Compared to LNCaP cells, C4-2B cells showed lesser or no repression of AR or AR target genes in response to TNFα (Fig. 4C & D); however, C4-2B cells did have an HS-5 CM response comparable to LNCaP cells (Fig. 3C & E). The respective sensitivity of LNCaP, LNas1, LNbs1 and C4-2B cells to IL-1, TNFα or HS-5 CM repression of AR and AR target genes is in line with their respective sensitivity to IL-1, TNFα or HS-5 CM cytotoxicity (Fig. 2A, 3B & 4A), where LNas1 and LNbs1 cells are the least sensitive. Thus, taken together, our data indicate that chronic IL-1 exposure selects for LNCaP cells that lose or attenuate intracellular responses to exogenous IL-1 or TNFα and further suggest that castration and chronic inflammation can both elicit similar loss or attenuation of intracellular inflammatory signaling.

### LNas1 and LNbs1 sublines have lower basal NFKB1/p50 levels

Canonical IL-1 signaling involves IL-1 binding to the IL-1R1 receptor leading to heterodimerization and nuclear translocation of the NFκB transactivating subunit, p65 (encoded by *RELA*), and the NFκB DNA-binding subunit, p50 (encoded by *NFKB1*)[1, 31]. We previously found that, while LNCaP and C4-2B cells have comparable IL-1R1 and RELA/p65 levels, C4-2B cells have lower NFKB1/p50 levels than LNCaP cells[15], which might underlie C4-2B reduced IL-1 sensitivity. To gain insight into the regulation of NFκB signaling in LNas1 and LNbs1 cells we analyzed mRNA and/or protein accumulation of IL-1R1, RELA/p65 and NFKB1/p50 in LNCaP, LNas1 and LNbs1 cells treated with 25 ng/ml IL-1α or IL-1β for 3 days. RT-qPCR and/or western blot analysis showed that LNCaP, LNas1 and LNbs1 cells have comparable IL-1R1 and RELA/p65 levels (Fig. 2B & D). However, LNas1 and LNbs1 cells have lower basal NFKB1/p50 levels than LNCaP cells and the sublines do not upregulate NFKB1/p50 in response to IL-1 (Fig. 2B & D). As with canonical IL-1 signaling, TNFα binding to its TNFR1 receptor also leads to p65:p50 heterodimerization, p65:p50 nuclear translocation and transactivation of NFκB target genes[1, 31]. Thus, the lower NFKB1/p50 levels in LNas1 and LNbs1 cells may underlie both IL-1 and TNFα insensitivity.

### AR shows functional activity in the LNas1 and LNbs1 sublines

Following acute (e.g., days) IL-1 repression of AR, AR levels and activity re-emerge in chronically (e.g., weeks) treated LNCaP cells (Fig. 1). In kind, the LNas1 and LNbs1 sublines have AR mRNA and protein levels comparable to LNCaP cells and show comparable or elevated mRNA and/or protein levels of AR target genes, *KLK2* and *NKX3.1* (Fig. 2B & D). To demonstrate that AR is active in LNas1 and LNbs1 cells, we siRNA-silenced *AR* (Fig. 5A & C) or, conversely, treated cells with the androgen analog, R1881 (Fig. 5B & D), and after 4 days of treatment, we determined the effect on KLK2 and NKX3.1 mRNA and/or protein levels by RT-qPCR and/or western blot. *AR* siRNA silencing reduced *KLK2* mRNA levels ∼50% in LNCaP, LNas1 and LNbs1 cells (Fig. 5A). While our RT-qPCR detected only a slight reduction on *NKX3.1* mRNA levels in LNCaP and LNbs1 (Fig. 5A), western blot analysis showed a decrease in NKX3.1 protein accumulation in LNCaP, LNas1 and LNbs1 cells transfected with *AR* siRNA (Fig. 5C). Conversely, 10 nM R1881 increased *KLK2* and *NKX3.1* mRNA (Fig. 5B) and NKX3.1 protein levels (Fig. 5D) in LNCaP, LNas1 and LNbs1 cells. Thus, as expected for the LNCaP cells, AR shows functional activity in the LNas1 and LNbs1 sublines as well.

**Fig 5.**
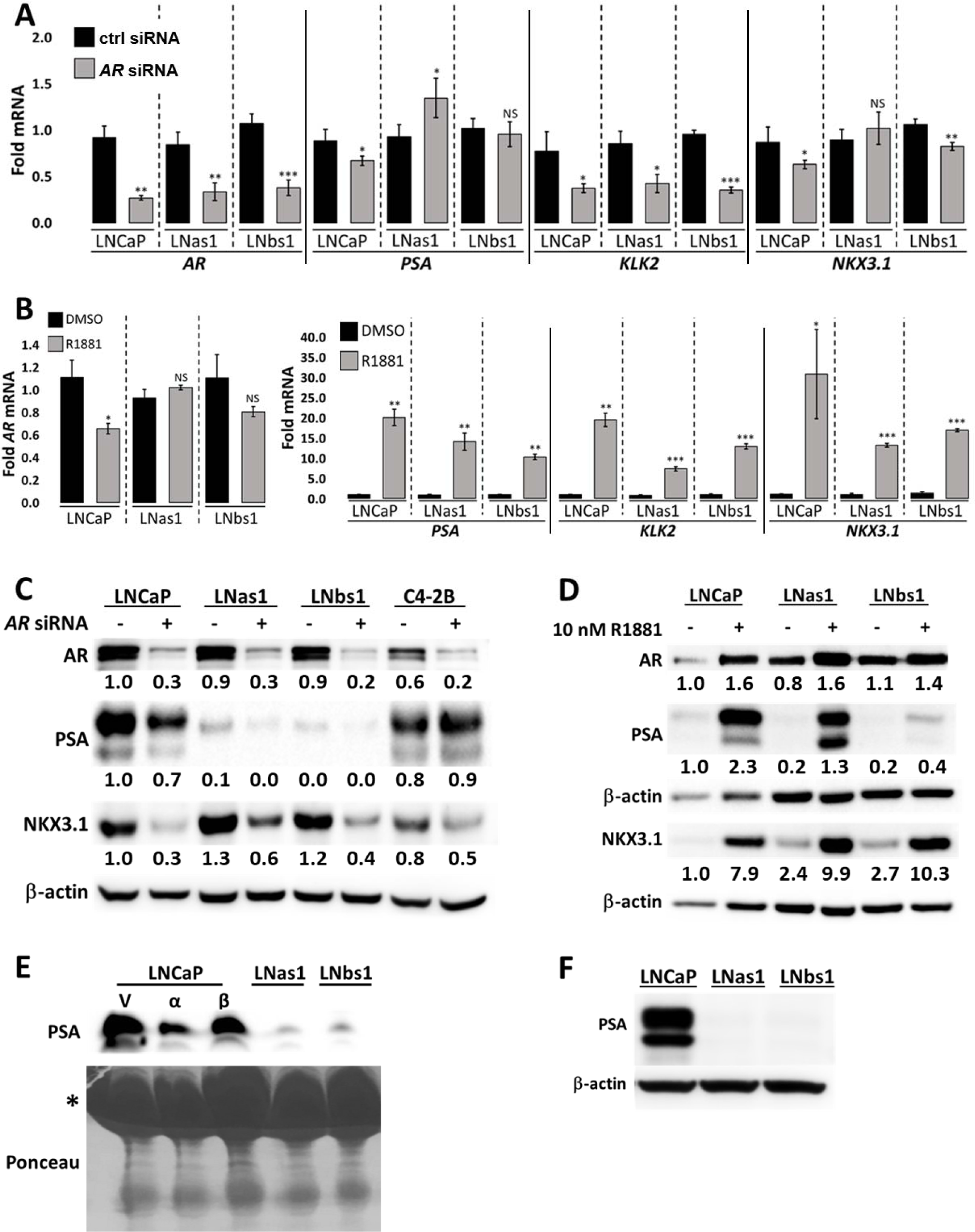
LNas1 and LNbs1 IL-1 sublines have active AR signaling but constitutively low PSA levels. LNCaP, LNas1, LNbs1 and C4-2B cells were transfected with 70 nM *AR* siRNA or siRNA control in DMEM/2.5% FBE or treated with 10 nM R1881 or vehicle control in DMEM/0% FBE and analyzed on day 4 for (A, B) RNA levels by RT-qPCR or (C, D) protein accumulation by western blot. AR siRNA reduced and R1881 increased the mRNA and/or protein accumulation of AR target genes *PSA*, *KLK2* and *NKX3.1* in LNCaP, LNas1 and LNbs1 cells. (E) LNCaP cells were treated with 25 ng/ml IL-1α or IL-1β for 3 days and LNas1 and LNbs1 cells were left untreated. Conditioned medium from the LNCaP, LNas1 and LNbs1 cells was analyzed for secreted PSA by western blot. LNCaP cells secreted PSA and IL-1 treatment reduced PSA secretion in LNCaP cells. LNas1 and LNbs1 secreted low levels of PSA compared to LNCaP cells. * indicates BSA on the ponceau stain. (F) LNCaP, LNas1 and LNbs1 were grown in DMEM/10% serum for 1 month and analyzed for PSA protein accumulation. PSA levels remain repressed in LNas1 and LNbs1 cells. Error bars, ± STDEV of 3 biological 0.05, ** 0.005, ***≤0.0005. For each individual cell line, fold mRNA levels are normalized to the cell line vehicle control; basal levels are not compared between the cell lines. Western blot band densitometry is normalized to β-actin and fold change normalized to LNCaP vehicle control.

### LNas1 and LNbs1 sublines show stable repression of *PSA* mRNA levels despite AR activity

Serum PSA is the surrogate biomarker for AR activity in PCa patients[14, 34]. Interestingly, while LNas1 and LNbs1 have active AR (Fig. 5), LNas1 and LNbs1 sublines have low or no detectable PSA (encoded by *KLK3*) mRNA or protein accumulation (Fig. 2B & D) or secreted PSA (Fig. 5E). Notably, although we did not detect a decrease in *PSA* mRNA levels in LNas1 or LNbs1 cells transfected with *AR* siRNA (Fig. 5A), we did detect a decrease in PSA protein (Fig.5C) and R1881 induced an increase in PSA mRNA and protein in LNCaP, LNas1 and LNbs1 cells (Fig. 5B & D). Thus, despite the low basal PSA levels in LNas1 and LNbs1, the AR activity in LNas1 and LNbs1 cells is still sufficient to regulate PSA levels.

To determine if the repression of PSA levels is transient or stable in the IL-1 sublines, we cultured the LNas1 and LNbs1 cells in normal growth medium with 10% serum for 1 month and we found that PSA accumulation remained repressed (Fig. 5F). Thus, the IL-1 sublines have evolved molecular changes that led to stable abrogation of canonical *PSA* expression. Taken together, while LNas1 and LNbs1 cells have active AR, some AR genes are mis-regulated.

### Differential gene expression analysis reveals the loss of canonical AR gene regulation in LNas1 and LNbs1 cells

Given that LNas1 and LNbs1 cells show constitutive repression of the AR target gene, *PSA/KLK3*, despite harboring functional AR activity (Fig. 2 & 5), we performed RNA sequencing of untreated LNCaP, LNas1 and LNbs1 cells to assess global changes in basal mRNA levels among the cell lines (GSE142706) (S1 Table 1). Differential gene expression analysis of basal mRNA levels in LNCaP, LNas1 and LNbs1 cells revealed that 2954 genes have higher basal expression in LNas1 and LNbs1 cells than LNCaP cells and 629 genes have lower basal expression (S1 Table 1).

Given that *PSA/KLK3* is constitutively repressed in LNas1 and LNbs1, we compared our 3583 gene set to a published gene set of R1881-induced genes in LNCaP cells[35] (GSE128749) to determine what other known R1881-induced genes are constitutively low in LNas1 and LNbs1 cells (S1 Table 1). Of the genes Poluri et al., found to be induced in LNCaP cells treated with 10 nM R1881 for 24 hours [35], 69 genes show basally low expression in LNas1 and LNbs1 cells, including *PSA/KLK3* (S1 Table 1). Conversely, of the genes that are induced by R1881 activity in LNCaP cells, 412 genes show basally high expression in LNas1 and LNbs1 cells, including *NKX3.1* and *KLK2* (S1 Table 1). In kind, there are genes repressed by R1881 that are basally low or high in the IL-1 sublines (S1 Table 1). Thus, while androgen/AR signaling is functional in LNas1 and LNbs1 cells (Fig. 5), global canonical AR regulation of gene expression is altered in LNas1 and LNbs1 cells, be it due to AR mis-regulation directly, or indirectly, due to other transcriptional and/or epigenetic changes.

### LNas1 and LNbs1 sublines have reduced dependency on AR for cell viability

Our data above indicates that chronic IL-1 exposure leads to direct or indirect molecular changes in canonical AR signaling. Therefore, we hypothesized that LNas1 and LNbs1 dependency on AR signaling for cell viability might also be altered. To test our hypothesis, we silenced *AR* using siRNA in LNCaP, LNas1, and LNbs1 cells and assayed cell viability using MTT (Fig. 6A). siRNA efficacy is shown for *AR* and AR target-gene mRNA and protein (Fig. 5A & C). siRNA-mediated loss of *AR* reduced cell viability ∼50% in LNCaP cells; however, LNas1 and LNbs1 cells showed little or no sensitivity to *AR* loss (Fig. 6A). Conversely, LNCaP, LNas1 and LNbs1 cells were grown in 10 nM R1881 in the absence of serum and assayed for cell viability to determine if androgen-induced activation of AR signaling improved cell viability in the IL-1 sublines (Fig. 6A). R1881 efficacy is shown for AR target-gene mRNA and protein (Fig. 5B & D). R1881 improved cell viability by ∼50% in LNCaP cells, by ∼20% in LNbs1 cells and had no significant effect on LNas1 cells (Fig. 6A). These data indicate that chronic IL-1 exposure selects for LNCaP cells that evolve reduced dependency on androgen and AR signaling for cell viability.

**Fig 6.**
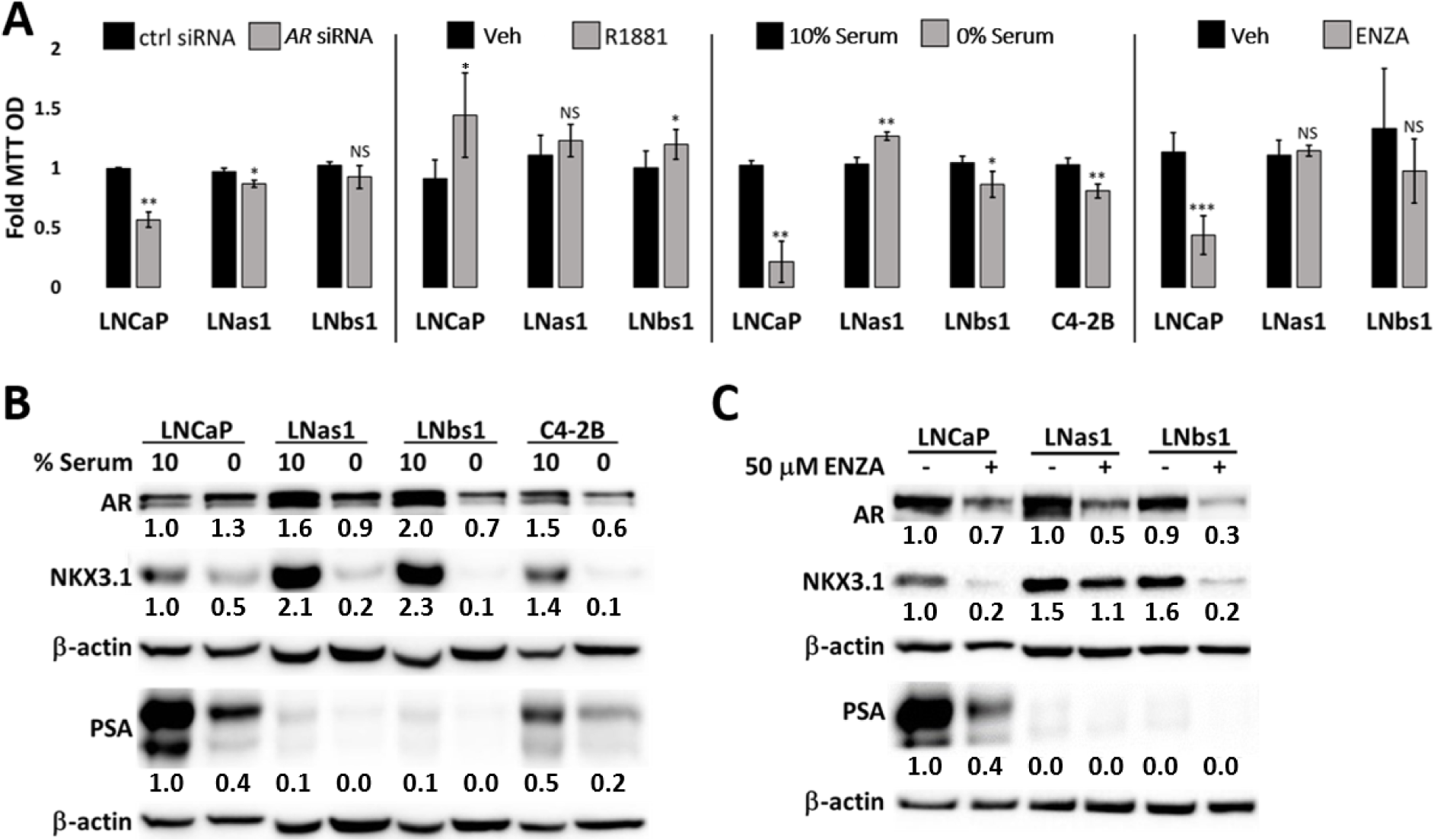
LNas1 and LNbs1 IL-1 sublines are insensitive to cytotoxicity induced by loss of AR or AR activity. (A) LNCaP, LNas1, LNbs1 and C4-2B cells were transfected with 70 nM *AR* siRNA or siRNA control in DMEM/2.5% FBE for 4 days, treated with 10 nM R1881 or vehicle control in DMEM/0% FBE for 4 days, grown in 10% versus 0% FBE (serum) for 5 days or treated with 50 μM enzalutamide (ENZA) for 5 days and analyzed for cell viability by MTT. LNas1, LNbs1 and C4-2B cells showed reduced sensitivity to *AR* siRNA, R1881, serum starvation or enzalutamide. (B) LNCaP, LNas1, LNbs1 and C4-2B cells were grown in 10% versus 0% FBE (serum) for 5 days and analyzed for protein accumulation of AR target genes, *NKX3.1* and *PSA*. Western blot shows that serum starvation (0% serum) reduced NKX3.1 and PSA accumulation, suggesting a reduction in AR activity. (C) LNCaP, LNas1 and LNbs1 cells were treated with 50 μM enzalutamide (ENZA) for 3 days and analyzed for protein accumulation of AR target genes, NKX3.1 and PSA. Western blot shows that enzalutamide reduced NKX3.1 and PSA accumulation. Error bars, ± STDEV of 3 biological replicates; p-value, * 0.05, ≤0.005, ***≤0.0005. For each individual cell line, fold MTT optical density (OD) is normalized to the vehicle control. Western blot band densitometry is normalized to β-actin and fold change normalized to LNCaP vehicle control.

### LNas1 and LNbs1 sublines have reduced dependency on serum for cell viability

The reduced dependency on androgen and AR-mediated cell viability in LNas1 and LNbs1 suggests that the sublines are also castration-resistant. Therefore, we grew LNCaP, LNas1 and LNbs1 cells in the presence (10% FBE) or absence (0% FBE) of serum for 5 days and determined cell viability using MTT (Fig. 6A). C4-2B cells are castration-resistant[36]; therefore, we used C4-2B cells as a treatment control. Serum starvation reduces AR activity; thus, for treatment efficacy, western blot shows that serum starvation reduced the protein accumulation of AR target genes, *PSA* and *NKX3.1* (Fig. 6B). Serum starvation reduced cell viability by ∼80% in LNCaP cells, by ∼15% in LNbs1 cells, by ∼20% in C4-2B cells and did not reduce cell viability in LNas1 cells (Fig. 6A). Thus, our data reveal that chronic IL-1 exposure selects for LNCaP cells that evolve reduced dependency on serum for cell viability.

### Chronic IL-1 exposure selects for LNCaP PCa cells that have reduced sensitivity to enzalutamide cytotoxicity

A second-generation therapy for CRPCa is the anti-androgen, enzalutamide, which binds AR to inhibit AR activity[37] and also reduces AR accumulation (Fig. 6C). LNCaP, LNas1 and LNbs1 cells were treated with 50 μM enzalutamide for 5 days and analyzed for cell viability by MTT (Fig. 6A). For treatment efficacy, western blot shows that enzalutamide reduced the protein accumulation of AR and AR target genes, *PSA* and *NKX3.1* (Fig. 6C). Enzalutamide reduced LNCaP cell viability by ∼60%, but LNas1 and LNbs1 showed no significant sensitivity to enzalutamide-induced cytotoxicity (Fig. 6A). Thus, our data reveal that chronic IL-1 exposure selects for LNCaP cells that evolved resistance to enzalutamide.

### Pathway analysis of differential gene expression data reveals the upregulation of known PCa pro-survival and pro-tumorigenic pathways in LNas1 and LNbs1 cells

Our data show that the IL-1 sublines co-evolved resistance to cytotoxic inflammation and the de-regulation of AR signaling. Thus, we hypothesized that the sublines upregulate other pro-survival, pro-tumorigenic pathways. IPA Canonical Pathway analysis of differentially expressed genes in untreated LNCaP versus untreated IL-1 sublines predicted that several pathways known to promote PCa survival, tumorigenicity or castration resistance are upregulated in LNas1 and LNbs1 cells (S1 Table 1). For example, EGF[38], AMPK[39], Wnt/Ca^2+^ [40], NGF[41], FGF[42], and ILK[43] signaling pathways promote PCa survival, tumorigenicity or castration resistance and are predicted to be constitutively high in the IL-1 sublines as compared to LNCaP cells (z-score ≥ 3.0, -log p-value ≥ 1.33) (S1 Table 1) and RT-qPCR confirms high basal expression of genes that mediate one or more of these pathways (S1 Table

1; S2 Fig.). Our data suggest that chronic IL-1 exposure selects for cells that can elicit pro-survival and pro-tumorigenic pathways that could compensate for the loss of canonical AR signaling and impart cyto-protection from cytotoxic inflammatory cytokines.

## Discussion

IL-1 is associated with disease progression in PCa patients[23, 24] and IL-1 promotes PCa metastasis and bone colonization[23, 44]. IL-1 induces EMT transition, transdifferentiation, and metastasis[23,44–47] and supports tumor growth and metastasis by inducing autocrine and paracrine secretion of molecules such as VEGF, IL-6, and IL-8[48]. Finally, IL-1 levels are elevated in PCa patient serum[24] and correlate with advanced Gleason score in primary tumors[23], indicating that IL-1 is clinically significant. With all that is known about IL-1 function in cancer, the role of IL-1 in PCa castration and anti-androgen resistance remains an area to be fully appreciated and explored.

PCa tumor cells require AR transcriptional activity for survival and proliferation and, thus, PCa therapies block androgen production (androgen deprivation therapy, ADT) or directly inhibit AR activity (anti-androgens)[8]. Patient tumors that develop ADT resistance are classified as castration-resistant PCa (CRPCa)[14]. Furthermore, a subset of CRPCa patients have intrinsic resistance to anti-androgens and almost all responsive patients will eventually develop anti-androgen resistance[49]. Mutated, variant or overexpressed AR underlie CPRCa[8] or anti-androgen resistance[49]. However, xenograft models demonstrate that castration- and anti-androgen-resistant PCa tumors can have low or no AR accumulation or activity[27]. In addition, in the era of anti-androgen use, there has been an increase in patients with PCa tumors that have low/no AR activity[42].

We and others found that acute IL-1 exposure represses AR accumulation and activity in AR- dependent PCa cell lines[4–7]; thus, we hypothesized that IL-1 selects for AR^low/-^ PCa cell populations that, consequently, acquire androgen and AR independence. But while acute inflammation is eventually resolved as part of host defense, unresolved chronic inflammation drives PCa initiation and progression[3]. Thus, we set out to determine if, in kind, chronic IL-1 exposure selects for AR^low/-^ PCa cell populations that, consequently, acquire androgen and AR independence. Staverosky et al. previously showed that acute IL-1β represses AR accumulation in LNCaP cells; but following a 3 week chronic IL-1β exposure, AR accumulation re-emerged and the LNCaP cells showed resistance to the anti-androgen, bicalutamide, *in vitro*[6]. We also found that within 3 weeks chronic IL-1α or IL-1β exposure, AR accumulation re-emerged in LNCaP cells (Fig. 1), but, in addition, we found that after 3-4 months’ exposure to IL-1α or IL-1β, LNCaP cells evolve aberrant canonical AR signaling (Fig. 2, S1 Table 1). For example, *PSA* expression is constitutively repressed, while *KLK2* and *NKX3.1* basal expression is elevated (Fig. 2, S1 Table 1). Furthermore, we discovered that LNCaP cells exposed to IL-1α or IL-1β for 3-4 months evolved resistance to serum starvation, *AR* silencing and enzalutamide (Fig. 5), suggesting that the cells are androgen- and AR-independent. Thus, while chronic IL-1 exposure did not select for AR^low/-^ LNCaP cells, as we had predicted it would be based on acute IL-1 repression of *AR* expression[4–7], chronic IL-1 exposure selected for CRPCa cells with aberrant canonical AR signaling (Fig. 2, S1 Table 1). Interestingly, while AR^-^ PCa cell lines, DU145 and PC3, do show IL-1 intracellular signaling responses, like LNas1 and LNbs1, the AR- cell lines are insensitive to IL-1-induced cytotoxicity (S3 Fig.). Thus, resistance to IL-1-induced cytotoxicity may reflect a progressive and aggressive PCa phenotype.

As shown, chronic IL-1 exposure led to stable repression of *PSA* (Fig. 5F). Notably, *PSA* expression can be regulated independent of AR[50–52], but it is unclear how chronic IL-1 exposure leads to stable *PSA* repression in the IL-1 sublines or mis-regulation of the other canonical AR target genes. Chronic IL-1 exposure may reorganize proximal and distal transcriptional regulators of *PSA* or other genes, thereby driving castration or anti-androgen resistance. Thus, it will be important to determine chronic IL-1 effect on chromatin remodeling and transcriptional regulation, including the regulation of the *PSA* loci and how this might affect CRPCa development.

In addition to the resistance to serum starvation, *AR* silencing, and enzalutamide, LNCaP cells evolved resistance to IL-1- and TNFα-induced cytotoxicity and intracellular signaling following 3- 4 months chronic IL-1 exposure (Fig. 2 & 4). IL-1 (*IL-1R1*, *IL-1RAcP*) and TNFα (*TNFR1*, *TNFR2*) receptor mRNA levels are comparable in LNCaP, LNas1 and LNbs1[15] (S1 Table 1), suggesting the evolved resistance occurs downstream of ligand-receptor interaction. NFκB mediates both IL-1 and TNFα intracellular signaling and canonical NF-κB signaling involves p65:p50 heterodimerization, p65:p50 nuclear translocation and transactivation of NFκB target genes[1, 31]. Thus, the lower basal p50 levels in LNas1 and LNbs1 cells (Fig. 2) may underlie both IL-1 and TNFα insensitivity. Mechanistic studies informed by sequencing and mass spectrometry data (data not shown) are underway to dissect how chronic IL-1 exposure leads to cytotoxic inflammatory cytokine insensitivity, as well as androgen and AR independence.

As part of the host defense response, IL-1 and TNFα increase reactive oxygen to induce cell death of invading pathogens[53, 54]. One way that cells can mitigate reactive oxygen stress is through the upregulation of antioxidant genes such as SOD2[55]. LNas1 and LNbs1, however, evolve resistance to the cytotoxic effects of IL-1 and TNFα and show no SOD2 induction in response to IL-1 or TNFα (Fig. 2 & 4). These results reflect the ability of cancer cells to co-opt immune responses to avoid cell death and promote tumorigenicity and, in response to chronic IL-1, develop NFκB-independent mechanisms to mitigate reactive oxygen stress. Indeed, canonical pathway analysis of differentially expressed genes in LNCaP versus the IL-1 sublines predicts the activation of multiple different pro-survival pathways in the IL-1 sublines (S1 Table 1) that might compensate for loss of canonical AR and NFκB signaling.

LNCaP cells were used to derive the castration-resistant isogenic cell line models, C4-2 and C4-2B, that, over a 16 week period of serial injections, were isolated from castrated mice co- inoculated with LNCaP and bone stromal cells[36]. Interestingly, the LNas1 and LNbs1 sublines that are derived from LNCaP cells exposed to chronic IL-1 phenocopy the C4-2 and C4-2B cells. C4-2[15], C4-2B[15], LNas1 and LNbs1 show reduced or no sensitivity to IL-1 (Fig. 2) and serum starvation (Fig. 6), have basally low or absent PSA and p50 accumulation and have elevated KLK2 and NKX3.1 levels (Fig. 2, S1 Table 1). Interestingly, Yu et al. observed tumor growth over a 6 week period using a LNCaP xenograft model and found that androgen deprivation induced PCa cell IL-1β secretion, leading to mesenchymal stem cell (MSC) recruitment, MSC CCL5 secretion and CCL5-dependent CRPCa development[12]. Furthermore, we have shown that bone marrow stromal cell-secreted IL-1 is cytotoxic for LNCaP cells but is less cytotoxic for C4-2B cells and has no significant effect on LNas1 or LNbs1 viability (Fig. 3). Thus, it is intriguing to speculate that C4-2 and C4-2B cell lines were derived from castration-resistant tumors that developed, at least in part, due to chronic IL-1 exposure in the tumor microenvironment.

Given that the tumor microenvironment is a complex community of multiple different cell types that interact to affect tumor cell behavior, including survival, proliferation, motility and autocrine and paracrine signaling, it will be important to investigate the LNas1 and LNbs1 sublines in the context of the tumor microenvironment to determine if chronic IL-1 exposure provides a growth advantage *in vivo*. While chronic IL-1 exposure renders the sublines resistant to IL-1- and TNFα-induced cytotoxicity, based on initial experiments (data not shown), we expect the LNas1 and LNbs1 sublines to remain responsive to growth factors and cytokines that support tumor progression, thereby conferring a growth advantage in the tumor microenvironment. In kind, Kawada and colleagues showed that an IL-1β-resistant LNCaP subline, generated by co-culturing LNCaP cells with a human fibroblast cell line in the presence of IL-1β, formed significantly larger xenografted tumors than parental LNCaP cells[56]. Importantly, the growth advantage rendered by chronic IL-1 exposure under castrate conditions and the paracrine influence of the sublines on the tumor microenvironment still need to be determined.

Our working model is that acute IL-1 exposure is largely cytotoxic at first for PCa cells, but concomitantly causes the emergence of an AR^low/-^ PCa subpopulation that is androgen- and AR- independent (Fig. 7). If the acute IL-1 inflammation is left unresolved, then chronic IL-1 selects from the AR^low/-^ PCa subpopulation for PCa cells that evolve resistance to IL-1 cytotoxicity, restore AR, and acquire reduced androgen and AR dependence (Fig. 7). While acute IL-1 responses can be mitigated with IL-1 antagonists, such as IL-1RA, to maintain or restore androgen and AR dependence and sensitivity to ADT and anti-androgens in PCa cells, chronic IL-1-induced phenotypes are not reversible and, therefore, require the identification of alternative therapeutic targets (Fig. 7).

**Fig 7.**
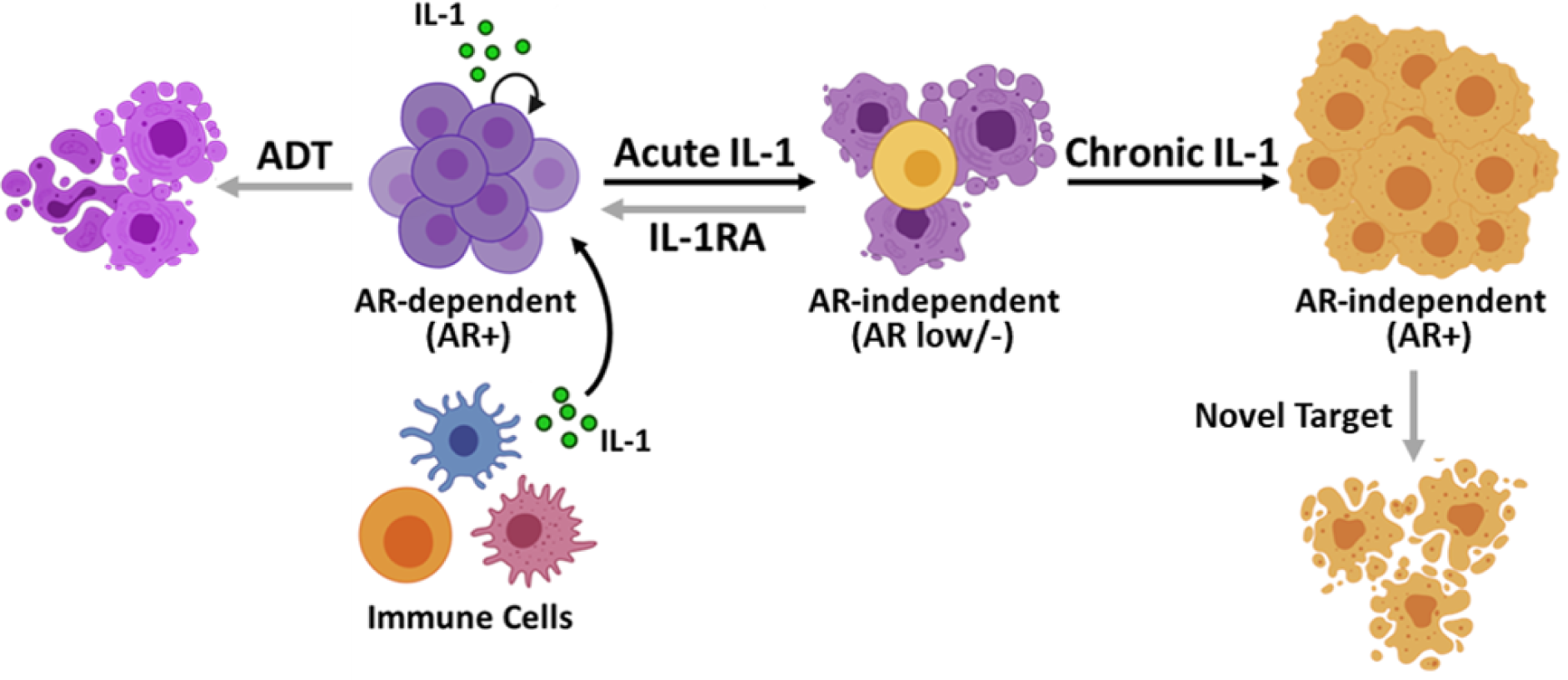
Model. IL-1 is secreted by immune and cancer cells in the tumor microenvironment. Acute IL-1 exposure is largely cytotoxic for PCa cells, but concomitantly causes the emergence of an AR^low/-^ PCa subpopulation that is androgen- and AR-independent. If left unresolved acute inflammation becomes chronic inflammation. Chronic IL-1 selects from the AR^low/-^ PCa subpopulation for PCa cells that evolve resistance to IL-1 (and TNFα) inflammatory cytokine cytotoxicity, restore AR, and acquire reduced androgen and AR dependence. Acute IL-1 responses can be mitigated with IL-1 antagonists, such as IL-1RA, to maintain or restore androgen and AR dependence and sensitivity to ADT and anti-androgens in PCa cells. Chronic IL-1-induced phenotypes are not reversible and, therefore, require the identification of alternative therapeutic targets.

## Conclusion

Acute inflammation is associated with a low risk of PCa development[57], which is in line the acute IL-1 cytotoxicity observed for LNCaP cells. But the clinical significance of chronic inflammation is unresolved, with studies showing association with increased[58–60] or, conversely, reduced PCa incidence[57, 61]. Irrespective, we show that, mechanistically, IL-1 can promote aggressive PCa phenotypes. Our data show that chronic IL-1 exposure can alter canonical AR signaling and reduce AR and androgen dependence in PCa cells. As such, the LNas1 and LNbs1 sublines are a power tool to investigate molecular (e.g., gene expression, epigenetics) or metabolic signatures that identify high risk PCa patients.

Elevated IL-1 accumulation is clinically significant in PCa progression and IL-1 has been shown to functionally promote PCa tumorigenicity, including the recruitment of mesenchymal stem cells to the primary tumor to promote CRPCa, metastasis and bone colonization. Importantly, CRPCa patients often develop androgen independent bone metastases. Therefore, chronic IL-1 exposure may contribute to CRPCa development in both the primary and metastatic tumor microenvironments.

Finally, intriguingly, we found that chronic IL-1 exposure can cause stable repression of the AR target gene, *PSA*. Serum PSA is the surrogate biomarker for AR activity in PCa patients. Therefore, our data emphasizes the importance of developing additional PCa biomarkers. A recent retrospective study conducted by Mahal et al. reveals that PSA low patients with high grade PCa have a poor prognosis[62]. Thus, it will also be important to determine if AR^+^/PSA^low/-^ PCa patients are predicted to develop CRPCa, show evidence of chronic IL-1- mediated tumor inflammation, and share a common molecular signature with the IL-1 sublines. Taken together, the IL-1 sublines are a powerful tool to identify alternative therapeutic targets for CRPCa.

## Acknowledgments

For their advice and support throughout this process, we would like to thank all the members of the Delk, Xing and Frigo labs. We would like to acknowledge the Genome Center at the University of Texas at Dallas and the DNA Genotyping Core at the University of Texas Southwestern Medical Center. We would also like to acknowledge financial support from the University of Texas at Dallas (Delk), NIH UL1TR001105 (Xing), NIH R01CA184208 and Prostate Cancer SPORE P50CA140388 (Frigo), and Antje Wuelfrath Gee and Harry Gee, Jr. Family Legacy Scholarship and Robert Hazelwood Graduate Fellowship for Cancer Research (Lin). The funders had no role in study design, data collection and analysis, decision to publish, or preparation of the manuscript.

## Author Contributions

HD, experiment design, execution, analysis, manuscript preparation. MK, bioinformatics analysis, manuscript preparation. STJ, optimization of cell culture, RT-qPCR, western blot analysis, manuscript editing. AN, cell survival and intracellular signaling experiments. SA and KP, undergraduate student researchers assisted HD with experiments. DF and CL assisted with data analysis. CX, head of bioinformatics core. ND, senior and corresponding author. All authors read and approved the final manuscript.

---

**S1 Table 1. Differential Gene Expression List and Pathway Analysis.** Differentially expressed genes (DEG) are listed for untreated LNCaP versus untreated LNas1 and LNbs1 (Tab 1_Subline DEG) and R1881-treated LNCaP cells (Tab 2_R1881 DEG). Pathway analysis was performed for LNCaP versus subline DEG (Tab 3_Subline Pathways). Finally, overlap (“venn”) in DEG lists for sublines and R1881 regulated genes are provided for the following: 1) R1881 upregulated and constitutively repressed in the sublines, 2) R1881 upregulated and constitutively upregulated in the sublines, 3) R1881 downregulated and constitutively repressed in the sublines, and 4) R1881 downregulated and constitutively upregulated in the sublines (Tab 4_Subline & R1881 Venn).

**S1 Figure.**
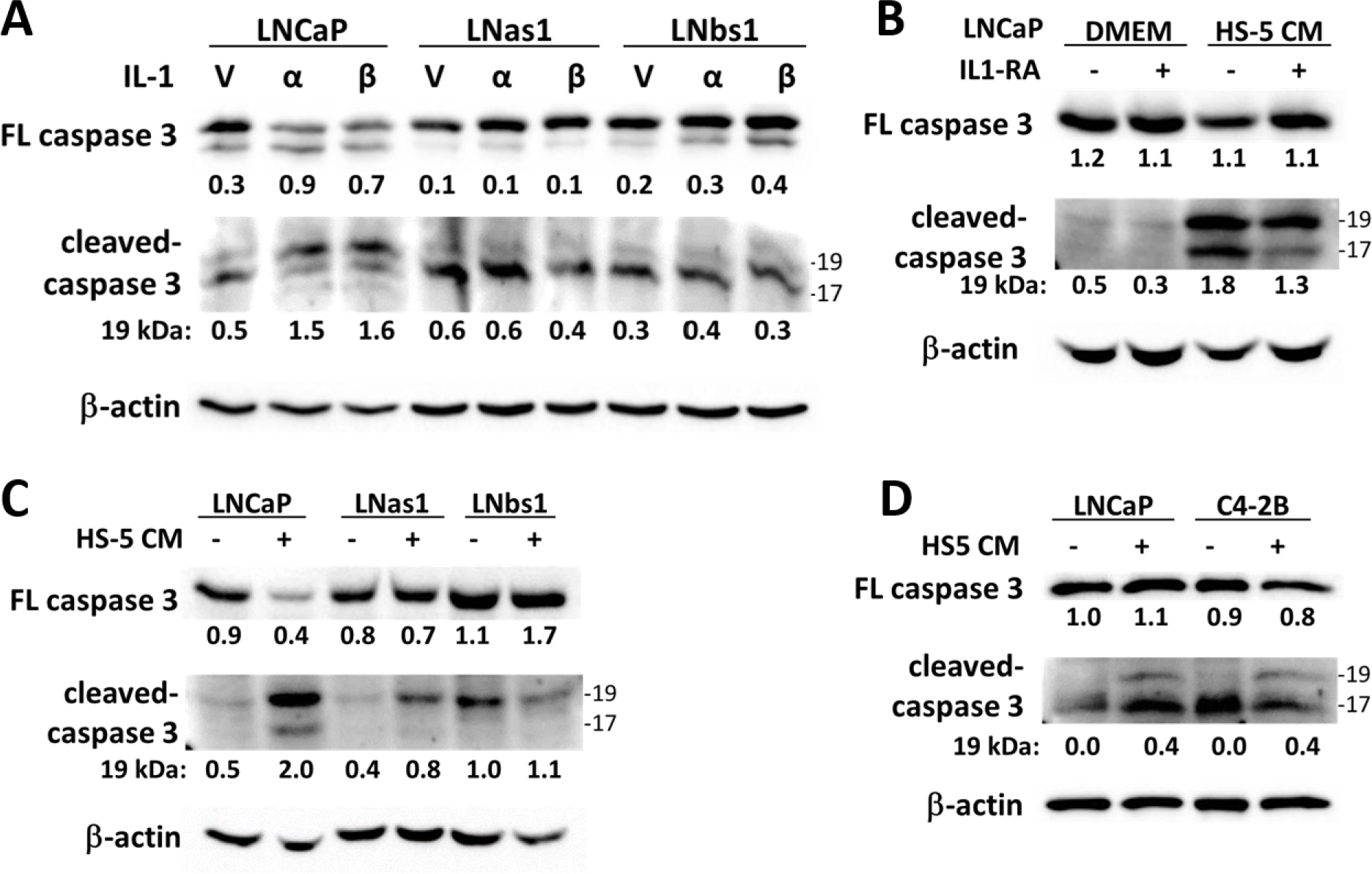
LNas1 and LNbs1 IL-1 sublines are resistant to IL-1-induced cytotoxicity (caspase 3 cleavage). (A) LNCaP, LNas1 and LNbs1 cells were treated for 3 days with vehicle control or 25 ng/ml IL-1α or IL-1β and analyzed for full-length (FL) caspase 3 protein accumulation using a caspase 3 antibody or caspase 3 low molecular weight cleavages products using a cleaved caspase 3 antibody. Detection of full-length caspase 3 turnover or cleavage products indicates activation of apoptosis. IL-1 induced cleavage of full-length caspase 3 and induced the accumulation of the 19 KDa caspase 3 cleavage product in LNCaP, but not LNas1 or LNbs1 cells. (B) LNCaP cells were pre-treated for 1 day with vehicle control or 400 ng/ml human recombinant IL-1RA and the following day the medium was replaced with D treatment control (DMEM) or HS-5 conditioned medium (CM) plus an additional 400 ng/ml D IL-1RA or vehicle control for 3 additional days. HS-5 CM reduced full-length caspase 3 and induced the accumulation of the 19 KDa and 17 KDa caspase 3 cleavage products. IL-1RA attenuated accumulation of the caspase 3 cleavage products. (C, D) LNCaP, LNas1, LNbs1 and C4-2B cells were treated for 3 days with treatment control or HS-5 CM. HS-5 CM reduced the accumulation of full-length caspase 3 and/or induced the accumulation of the 19 KDa and/or 17 KDa cleavage products in LNCaP and C4-2B cells, but not in LNas1 or LNbs1. Taken together, LNas1 and LNbs1 have reduced sensitivity to IL-1-induced apoptosis activation. Full-length caspase 3 cleavage densitometry shows the ratio of cleaved to uncleaved caspase 3. Low molecular weight 19 KDa and 17 KDa caspase 3 cleavage product densitometry is normalized to β-actin.

**S2 Figure.**
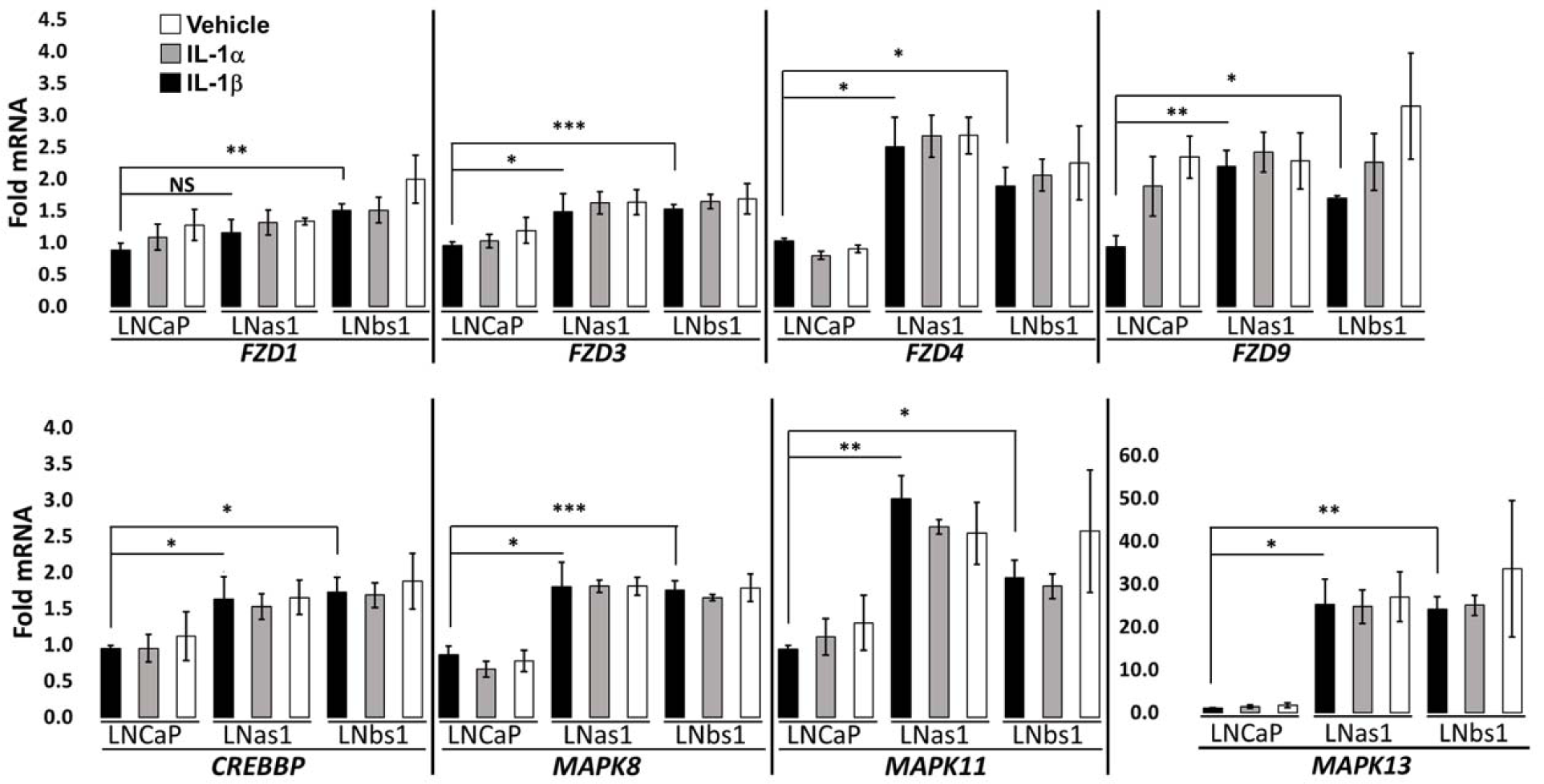
LNas1 and LNbs1 IL-1 sublines show high basal expression of genes that mediate pathways known to promote PCa survival, tumorigenicity or castration resistance. LNCaP, LNas1 and LNbs1 cells were treated for 3 days with vehicle control or 25 ng/ml IL-1α or IL-1β and analyzed for mRNA levels by RT-qPCR for *FDZ1*, *FDZ3*, *FDZ4*, *FDZ9*, *CREBBP*, *MAPK8*, *MAPK11*, *MAPK13*. These genes were chosen arbitrarily from S1 Table 1 IPA analysis to represent a cross section of the EGF, AMPK, Wnt/Ca2+, NGF, FGF and ILK pathways. While acute IL-1 exposure did not modulate the expression level of the genes in LNCaP, LNas1 or LNbs1, basal gene expression was high in LNas1 and LNbs1. Error bars, ± 0.05, ** 0.005, *** 0.0005, NS = not significant. ≤ Fold mRNA levels are normalized to LNCaP vehicle control for IL-1 treatments in order to also compare basal levels between the cell lines.

**S3 Figure.**
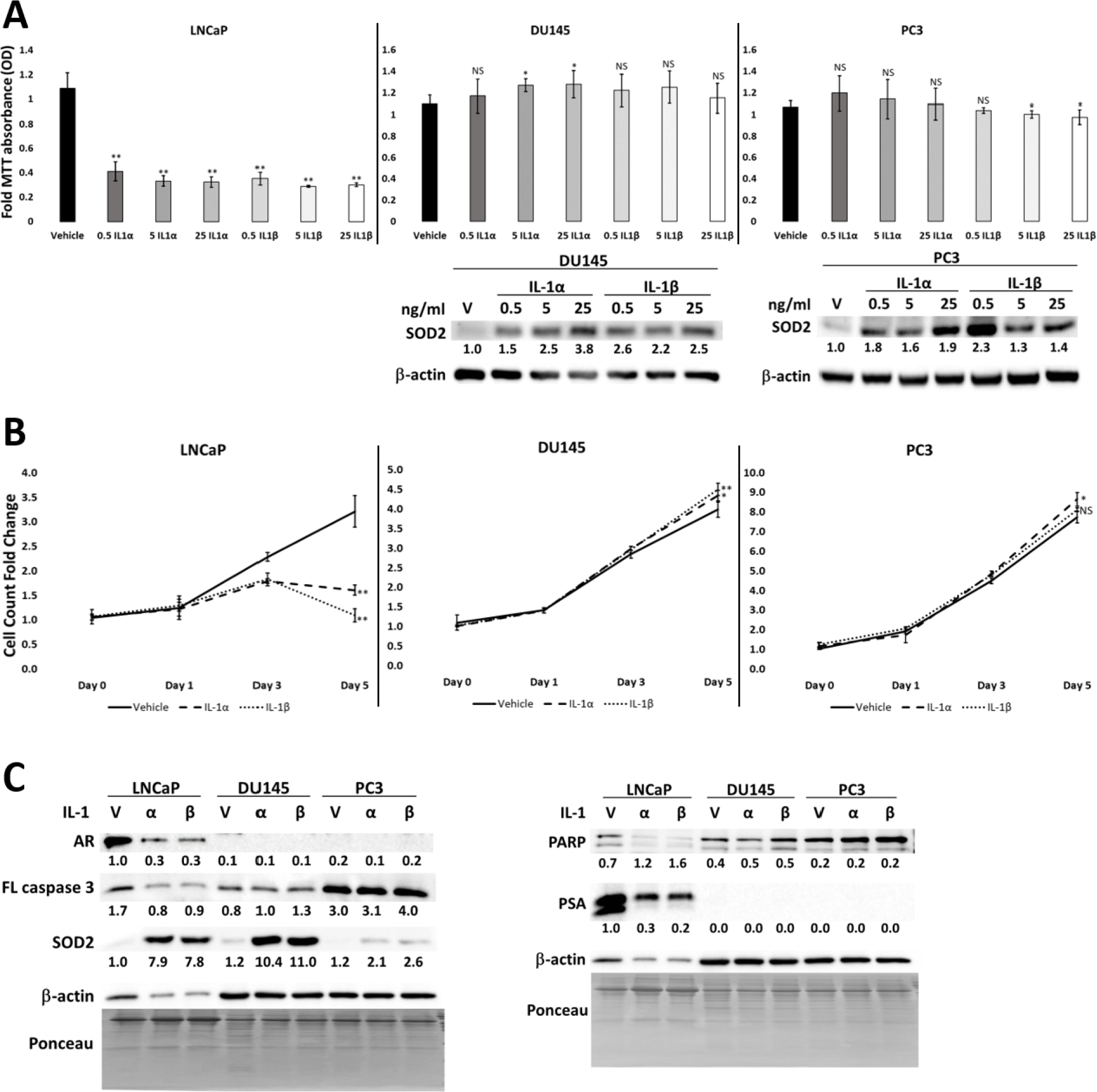
AR-negative PCa cells lines are insensitive to IL-1-induced cytotoxicity. (A) LNCaP, DU145 and PC3 cells were treated for 3 days with vehicle control or 0.5-25 ng/ml IL-1α or IL-1β and analyzed for cell viability using MTT or for IL-1-induced intracellular signaling using SOD2 protein accumulation. LNCaP cells are sensitive to IL-1 induced cytotoxicity. No cells remained for protein analysis. DU145 and PC3 cells activated IL-1 intracellular signaling, but were not sensitive to IL-1-induced cytotoxicity. (B) LNCaP, DU145 and PC3 cells were treated for 0-5 days with vehicle control or 25 ng/ml IL-1α or IL-1β and viable cell counts determined. IL- 1 reduced LNCaP cell number over time, but had no appreciable effect on DU145 or PC3 cell counts. (C) LNCaP, DU145 and PC3 cells were treated for 5 days with vehicle control or 25 ng/ml IL-1α or IL-1β and analyzed for AR, PSA, SOD2, PARP cleavage, and full-length caspase 3. As expected, IL-1 reduced AR, PSA and full-length caspase 3 accumulation and induced SOD2 accumulation and PARP cleavage in LNCaP cells. AR-negative DU145 and PC3 cell lines showed IL-1-induced SOD2 accumulation, but did not show IL-1-induced caspase 3 turnover or PARP cleavage. Error bars, ± STDEV of 3 biological replicates; p-value, * 0.05, ≤ ** 0.005, NS = not significant. Fold MTT optical density (OD) and cell counts are normalized to ≤ the treatment control. Low β-actin in IL-1-treated LNCaP cells reflects cell death; therefore, protein bands for LNCaP, DU145 and PC3 were normalized to ponceau for densitometry and PARP densitometry shows the ratio of cleaved to uncleaved PARP.

